# Quantifying nucleation *in vivo* reveals the physical basis of prion-like phase behavior

**DOI:** 10.1101/205690

**Authors:** Tarique Khan, Tejbir S. Kandola, Jianzheng Wu, Shriram Venkatesan, Ellen Ketter, Jeffrey J. Lange, Alejandro Rodríguez Gama, Andrew Box, Jay R. Unruh, Malcolm Cook, Randal Halfmann

**Author notes:** These authors contributed equally (Tarique Khan, Tejbir S. Kandola, Jianzheng Wu, Shriram Venkatesan, Ellen Ketter).

## Abstract

Protein self-assemblies modulate protein activities over biological time scales that can exceed the lifetimes of the proteins or even the cells that harbor them. We hypothesized that these time scales relate to kinetic barriers inherent to the nucleation of ordered phases. To investigate nucleation barriers in living cells, we developed Distributed Amphifluoric FRET (DAmFRET). DAmFRET exploits a photoconvertible fluorophore, heterogeneous expression, and large cell numbers to quantify via flow cytometry the extent of a protein’s self-assembly as a function of cellular concentration. We show that kinetic barriers limit the nucleation of ordered self-assemblies, and that the persistence of the barriers with respect to concentration relates to structure. Supersaturation resulting from sequence-encoded nucleation barriers gave rise to prion behavior, and enabled a prion-forming protein, Sup35 PrD, to partition into dynamic intracellular condensates or to form toxic aggregates. Our results suggest that nucleation barriers govern cytoplasmic inheritance, subcellular organization, and proteotoxicity.

**Highlights:**

- Distributed Amphifluoric FRET (DAmFRET) quantifies nucleation in living cells
- DAmFRET rapidly distinguishes prion-like from non-prion phase transitions
- Nucleation barriers allow switch-like temporal control of protein activity
- Sequence-intrinsic features determine the concentration-dependence of nucleation barriers

## INTRODUCTION

Ordered protein self assemblies function to transduce biological signals and encode molecular memories (Banani et al., 2017; Caudron and Barral, 2013; Chakrabortee et al., 2016; Halfmann, 2016; Si and Kandel, 2016; Wu and Fuxreiter, 2016), but they also precipitate incurable degenerative diseases like Alzheimer’s, Parkinson’s, and ALS (reviewed in Knowles et al., 2014). These phenomena emerge over much longer timescales than those typical either of protein folding or of liquid-liquid phase separation.

At the extreme, remarkable self-assemblies known as prions govern protein activity over timescales that can span multiple generations of the host organism. Prion-forming proteins normally exist as dispersed monomers. But this state is only kinetically stable with respect to a thermodynamically-favored, assembled state (Glover et al., 1997; Tanaka et al., 2006). The assemblies typically take the form of exquisitely ordered quasi-two-dimensional polymers known as amyloids (Kashchiev, 2015; Kashchiev and Auer, 2010; Nelson et al., 2005; Tycko and Wickner, 2013; Wasmer et al., 2008; Zhang and Schmit, 2016; Zhao and Moore, 2003). Prion assemblies appear spontaneously at very low frequencies, but can be “induced” by transient over-expression of the protein. Once formed, they template the conversion of other molecules of the protein to the same state (Prusiner, 1982; Satpute-Krishnan and Serio, 2005; Tanaka et al., 2006). Should a fragment of the assembly then enter a naive pool of the protein within a foreign cell or organism, it converts them as well. This capability grants prions properties otherwise found only in nucleic acids – the ability to transmit phenotypes between organisms and across generations.

The many molecular degrees of freedom that must be lost upon ordered self-assembly *de novo* can render that event, known as nucleation, inherently probabilistic on the molecular scale. The “nucleation barrier” describes the extent of that improbability. It therefore theoretically determines the fraction of otherwise identical systems that will acquire the assembly spontaneously over a given period of time, as well as how long, on average, any single system will remain free of it. Although nucleation barriers are well-grounded in theory (Michaels et al., 2017; Vekilov, 2012; ten Wolde and Frenkel, 1997), their relevance to complex biological phenomena is underexplored.

To what extent do nucleation barriers govern protein kinetics in biological systems? Quantifying low probability nucleation events under cellular conditions is crucial to answering this question. We developed a facile approach – DAmFRET – to do so. DAmFRET measures the frequency of nucleation as a function of protein concentration in living cells. Applying it to diverse proteins, we reveal that sequence-encoded nucleation barriers relate to the structures and functions of self-assemblies. Our findings suggest that nucleation barriers broadly govern supersaturation-dependent protein activities, ranging from prion behavior, to stress-responsive condensation, to proteotoxicity.

## RESULTS

### Distributed Amphifluoric FRET (DAmFRET) reveals nucleation barriers to self-assembly in the cellular milieu

Self-assemblies differ from their corresponding unassembled polypeptides in one or more critical order parameters. These include density, and – depending on the assembly – orientation and conformation. Nucleation occurs when random fluctuations in those parameters happen to produce a minimal cluster of the polypeptides thermodynamically sufficient for further growth (Cellmer et al., 2016; Kashchiev and Auer, 2010; Powers and Powers, 2006; Vekilov, 2012). In sufficiently small systems, the improbability of those fluctuations occurring simultaneously results in a kinetic barrier that can allow soluble proteins to accumulate beyond the minimal – or “saturating” – concentration required for growth. Such supersaturated systems go on to nucleate in probabilistic fashion, resulting in a collective bimodal dependence of assembly on concentration.

Detecting nucleation barriers therefore calls for the examination of very large numbers of independent molecular systems over a wide range of concentrations. Compartmentalizing purified proteins into microdroplets is one approach to do so (Ildefonso et al., 2012; Michaels et al., 2017; Peters, 2011). However, it divorces proteins from myriad intracellular factors that influence their structure. Living cells offer a more physiological approach.

We therefore sought a single-cell reporter of protein self-assembly which could be assessed in large populations of independent cells expressing the protein over a range of concentrations. To achieve this goal, the system would have to: provide single-cell readouts; scale to report on thousands of cells in a population; ensure independence between cells; be manipulable to produce expression over a wide concentration range; provide a sensitive readout of protein expression and cytosolic volume (as required for determining concentration); and work equally well across dozens of different target proteins.

The necessity to acquire single-cell measurements across large cell populations suggested flow cytometry as the appropriate platform. The need to determine cytosolic volume and the desire to evaluate protein localization within each cell further led us to imaging flow cytometry (Basiji and O’Gorman, 2015)

Ensuring experimental independence of each cell required that we restrict intercellular interactions. We therefore employed the unicellular eukaryote, budding yeast. Notably, yeast cell walls are impermeable to extracellular amyloids and prion-containing exosomes (Kabani and Melki, 2015; King and Diaz-Avalos, 2004) whereas cultured mammalian cells readily internalize these structures (Grassmann et al., 2013). We eliminated the additional possibility of mitotic inheritance of nucleated assemblies by genetically inducing cell cycle arrest during query protein expression (see Methods). In short, every cell of the resulting yeast strain can be considered as an independent, femtoliter-volume vessel for protein assembly.

Many techniques were considered for the reporter. Förster Resonance Energy Transfer (FRET) occurs between two fluorophores that have overlapping spectra, when they are brought in close proximity. FRET is widely used to detect interactions between two corresponding protein species fused to those fluorophores (Jares-Erijman and Jovin, 2003). However, creating the two fusions for each protein, and expressing them to the same ratio in all cells across a range of expression levels, would be exceedingly difficult. We likewise excluded protein complementation and two hybrid assays from consideration. Available approaches that use a single fusion protein to detect self-assembly, such as fluorescence anisotropy and enzyme loss of function assays, were not considered due to their limited dynamic range and throughput.

We reasoned that high-throughput, sensitized emission FRET between complementary fluorophores could be realized if they were both expressed from the same genetic construct. Individual molecules of the fusion protein would have to mature stochastically into one or the other fluorophore, resulting in a mixture of the two at the cellular level. In this way, a consistent ratio of donor to acceptor molecules could be produced regardless of the protein’s expression level. We further reasoned that photoconvertible fluorescent proteins could be employed for this purpose (Wolf et al., 2013). After testing several, we chose to proceed with mEos3.1, a monomeric bright green fluorescent protein that can be converted irreversibly to a bright red fluorescent form upon illumination with violet light (Zhang et al., 2012). Importantly, the emission spectrum of the green form strongly overlaps the excitation spectrum of the red form, as necessary for FRET to occur (Fig. 1A). The ratio of green to red molecules and thereby the sensitivity of the mixture to self-assembly can be precisely controlled by modulating the intensity and duration of violet light exposure. We term this approach <u>A</u>mphifluoric <u>FRET</u> (AmFRET) due to the dual nature of the fluorescent moiety.

**Figure 1.**
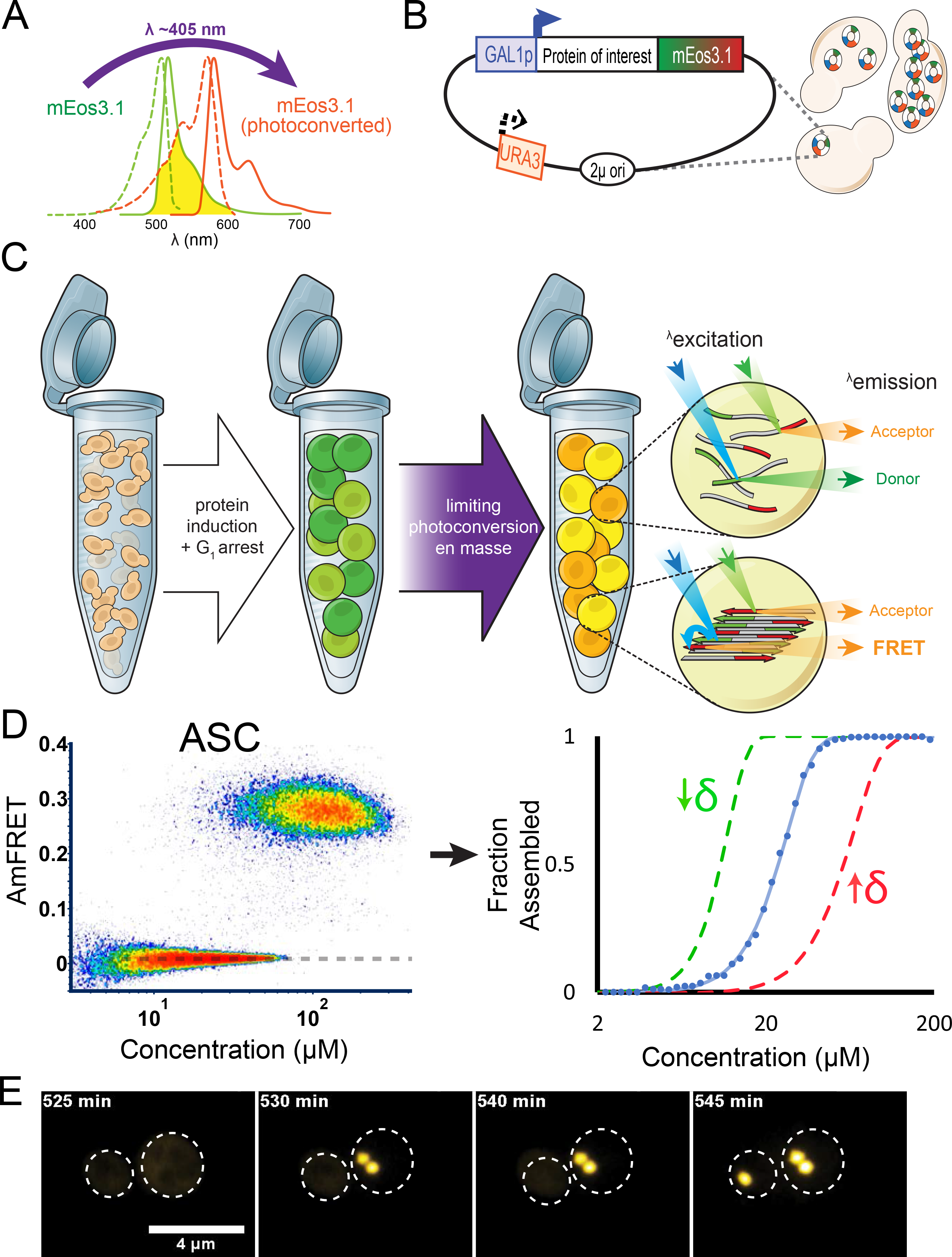
Distributed Amphifluoric FRET (DAmFRET) reveals nucleation barriers to self-assembly in the cellular milieu. (A) Violet light photoconverts mEos3.1 from a green to a red form. The emission (solid curves) and excitation (dotted curves) spectra for the green form and red form of mEos3.1 are shown in green and red curves, respectively; and the region of overlap between them, responsible for FRET, is shaded in yellow. (B) Schematic representation of the genetic constructs used for DAmFRET. The expression of the query protein-mEos3.1 fusion is driven by a strong inducible promoter (*GAL1*) from a plasmid which accumulates to highly variable copy numbers among yeast cells owing to its 2μ origin of replication and very low expression of the auxotrophic marker (*URA3*). (C) Cartoon depicting DAmFRET. A culture of cells harboring the query protein-mEos3.1 fusion construct (left tube), is shifted to medium that simultaneously induces query protein expression while arresting cell division, resulting in large, spherical cells that accumulate the query protein to a wide range of concentrations (middle tube). The culture of cells is then uniformly irradiated with violet light for a limited duration to convert an empirically optimized fraction of mEos3.1 molecules to the red form (right tube). When in close proximity to one another, red molecules will fluorescence upon excitation of green molecules (FRET). Non-colinearity of the 488 nm (in blue) and 561 nm (in green) lasers in the ImageStream^®x^ MkII ensures that direct and sensitized emission (FRET) of the red molecules can be distinguished. See also Fig. S1. (D) DAmFRET plot of human ASC. The dashed line approximates the mean AmFRET value of cells expressing fully monomeric protein. All DAmFRET plots represent at least 20,000 cells. The blue curve to the right represents the phenomenological fit of the DAmFRET plot to a Weibull distribution (see Methods). The parameter **δ** relates to the sharpness of the transition, and describes the persistence of the low population with respect to concentration. The green and red curves represent hypothetical distributions with values of **δ** that are lower or higher, respectively, than that of the blue curve. (E) Montage of cells expressing ASC protein, showing switch-like acquisition of puncta. Images represent sum projection of confocal slices captured at 63X magnification.

To detect nucleation barriers, we needed to evaluate the <u>d</u>istribution of <u>AmFRET</u> values, or DAmFRET, across a wide range of intracellular protein concentrations. We therefore inducibly expressed mEos3.1 fusion proteins from an episomal plasmid whose copy number varies more than one hundred fold between cells (Fig. 1B; Futcher and Cox, 1984; Loison et al., 1989). After eighteen hours of protein expression, cell cultures were uniformly illuminated with violet light to convert a fraction of mEos3.1 molecules from the green (donor) to the red (acceptor) form (Fig. 1C). We limited the illumination to an empirically optimized dose that yielded maximum FRET intensity. We established that photoconversion efficiency was insensitive to expression level as well as to the identity and structure of the fusion partner (Fig. S1B), and did not vary between cells (Fig. S1C), enabling us to indirectly measure total protein levels as the product of intensity of acceptor fluorescence and an empirically determined molecular brightness and photoconversion factor (see Methods). We then divided each cell’s total protein level by its approximate cytosolic volume as calculated from the bright-field image to derive absolute protein concentrations. We used the ratio of acceptor fluorescence when excited indirectly (at 488 nm) to directly (at 561 nm) to approximate FRET efficiency, and hereafter refer to this ratio as simply, “AmFRET”.

To validate DAmFRET, we used the well-characterized human signaling protein, ASC, whose nucleation into a right-handed triple helical polymer confers a digital, all-or-none responsiveness to inflammatory stimuli (Cai et al., 2014; Cheng et al., 2010; Lu et al., 2014). We had previously demonstrated that ASC acquires its physiological polymeric form when expressed in yeast (Cai et al., 2014).

We found that cells expressing ASC-mEos3.1 to low concentrations lacked AmFRET. But at higher concentrations, a second population emerged with intense AmFRET (Fig. 1D). The two populations were discontinuous yet overlapped on the abscissa, resulting in a strongly bimodal distribution at intermediate concentrations. This overlap indicates the existence of discrete phases of the protein, and that the transition from one to the other is not determined solely by concentration on the time scale of our experiment. The transition is therefore also subject to a kinetic barrier, which we attribute to nucleation. Cells expressing mEos3.1 without afusion partner produced negligible AmFRET even at the highest concentrations measured – approximately 200 μM (Fig. S1G).

To describe the observed persistence of the nucleation barrier with respect to concentration, we used a single parameter extracted from the data as follows. We gated the distribution into AmFRET-negative and AmFRET-positive cell populations, representing pre- and post-nucleation events, respectively. We fit the fraction of cells in the AmFRET-positive population to a Weibull distribution (Fig. 1D; see also Methods and Table S1), a simple and purely phenomenological model that has been used to describe nucleation probability as a function of supersaturation (Sear, 2016). The dimensionless Weibull shape parameter describes the sharpness of the transition (Rinne, 2008). We therefore use its reciprocal, designated **δ**, to report the observed persistence of the nucleation barrier with respect to concentration. The minimum theoretical value of **δ**, corresponding to self-assembly without a detectable nucleation barrier, is 0. There is no maximum value.

To further validate that overlapping AmFRET results from a nucleation barrier, we tested two additional predictions. 1) If the AmFRET-positive state of ASC is indeed rate-limited by nucleation, then on average, it will occur only once per cell. Absent secondary processes like fragmentation, this must result in a single fluorescent punctum. An analysis of the imaging data revealed that most cells in the top population indeed contained a single intensely fluorescent punctum from which the FRET signal originated (Fig. S1F). Cells in the bottom population contained fully dispersed fluorescence. 2) The absence of cells with intermediate values of AmFRET indicates that the single nucleus grows so rapidly as to achieve steady state near instantaneously. Otherwise, we would observe cells in transition between the lower and upper populations. To evaluate, we recorded the expression level and distribution of ASC over time in multiple individual yeast cells. We found that fluorescence accumulated to high levels in a fully diffuse state. But then, in a stochastic fashion for each cell, it collapsed near-instantaneously into discrete puncta (Movie S1 and Fig. 1E). These kinetics and puncta morphology closely resemble ASC activation in human cells (Cheng et al., 2010). We conclude that DAmFRET accurately reported a kinetic barrier attributable to polymer nucleation.

### Amyloid structure determines the persistence of the nucleation barrier with respect to concentration

The extent to which a protein’s structure changes upon assembly must, theoretically, contribute to the barrier to nucleating that assembly. Assemblies of different structure will therefore correspond to different AmFRET distributions.

To evaluate if **δ** indeed contains information about structure, we required a single protein that can variously nucleate into structurally distinct self-assembled forms. For this purpose we chose the prion-determining region (PrD) of the archetypal yeast prion protein, Sup35. Sup35 PrD is unstructured as a monomer (Mukhopadhyay et al., 2007), but forms a spectrum of mutually exclusive amyloid isoforms whose ordered cores incorporate different lengths of the polypeptide backbone (Tanaka et al., 2004). The distribution of isoforms is governed by another protein, Rnq1, whose [*PIN*+^high^], [*PIN*+^medium^], and [*PIN*+^low^] isoforms preferentially induce Sup35 amyloids with relatively short, medium, and long ordered cores, respectively (Bradley et al., 2002; Tanaka et al., 2004; Westergard and True, 2014). Sup35 PrD amyloids virtually never nucleate in cells with an inactive Rnq1 variant ([*pin-*]). Our hypothesis predicts that a) Sup35 PrD will produce bimodal distributions of cells with discrete populations corresponding to its non-prion and prion ([*PSI*+]) states, b) that the distribution of cells between the two populations will differ between Rnq1 variants, and most specifically, c) that **δ** of those distributions will increase from [*PIN*+^high^] to [*PIN*+^medium^] to [*PIN*+^low^] to [*pin-*].

We first verified that Sup35 PrD acquired a high AmFRET state in cells harboring prions of endogenous Sup35 ([*PSI+*]; Fig. 2A). We next asked if expression of Sup35 PrD in the absence of Sup35 prions produced bimodal DAmFRET, as anticipated for prion nucleation de novo. It did (Fig. 2B). To verify that these high AmFRET cells, specifically, contained prions, we employed a well-established genetic background that causes cells to accumulate red pigment when they contain soluble Sup35, but not when they contain aggregated Sup35 (Osherovich et al., 2004). We used fluorescence-activated cell sorting (FACS) to isolate AmFRET-positive and AmFRET-negative cells expressing the same amount of Sup35 PrD, and then plated them to media that repressed its expression. Cells in the lower population formed red colonies, whereas cells in the upper population formed a spectrum of white and pink colonies (Fig. 2C), demonstrating that the AmFRET-positive population contained independently nucleated self-perpetuating amyloid isoforms.

**Figure 2.**
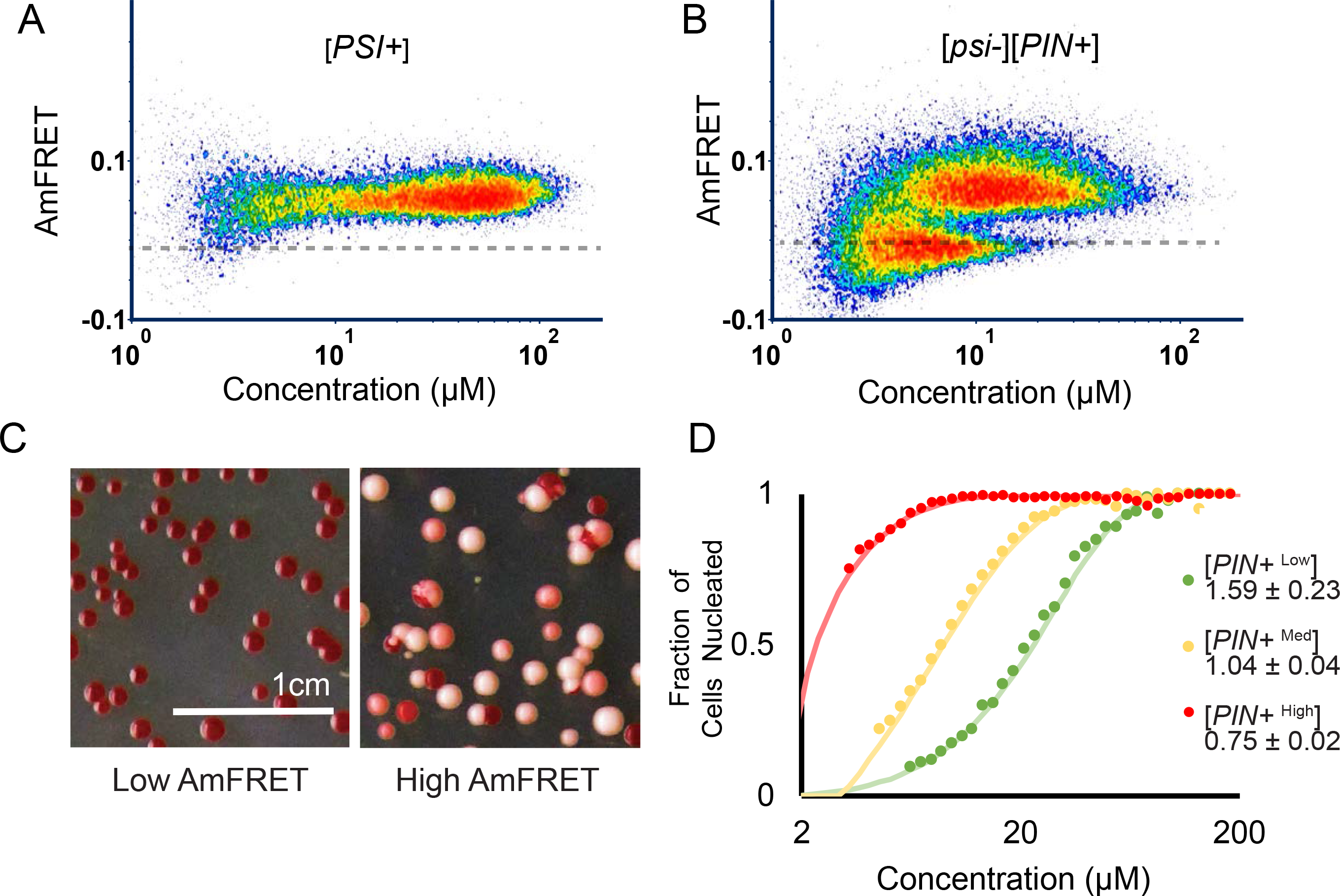
Amyloid structure determines the persistence of the nucleation barrier with respect to concentration. (A-B) DAmFRET plots of Sup35 PrD in cells with amyloids of endogenous Rnq1 ([*PIN*+]) and with (A; [*PSI*+]) or without (B; [*psi*-]) amyloids of endogenous Sup35. (C) Representative images of colonies of cells from Sup35 colony color assay. Left and right panels show colonies grown from cells sorted from the low or high AmFRET populations, respectively, as shown in figure 2B. (D) Weibull fits of DAmFRET of Sup35 PrD in cells with different amyloid isoforms of endogenous Rnq1:[*PIN*+^high^], [*PIN*+^medium^], and [*PIN*+^low^]. The inset shows **δ** ± error.

We then proceeded to acquire DAmFRET for Sup35 PrD in the presence of endogenous Rnq1 amyloid isoforms. Exactly as predicted, **δ** increased from [*PIN*+^high^] to [*PIN*+^medium^] to [*PIN*+^low^] (Fig. 2D, S2). [*pin-*] cells were entirely unable to populate the high AmFRET state, suggesting that Sup35 PrD sans conformational template supersaturates to a depth that exceeds the sensitivity of DAmFRET. We conclude that **δ** correlates with conformational ordering in amyloid isoforms.

### Nucleation barriers govern prion behavior

A prion only exists with respect to a metastable, non-prion state of the same protein. Otherwise the defining activity of prions – transmission – could not occur. We therefore submit that **δ** is the single most informative descriptor of prion behavior at the cellular level.

To evaluate, we expanded our analysis to multiple proteins that share Sup35 PrD’s characteristic low complexity and “prion-like” enrichment for polar, uncharged residues. These included PrDs from multiple other yeast prion proteins: Ure2, Rnq1, Swi1, Mot3, and Cyc8 (Alberti et al., 2009; Du et al., 2008; Patel et al., 2009; Sondheimer and Lindquist, 2000; Wickner, 1994), as well as the prion-like regions (PrL) of the yeast proteins Ngr1 and Sla1. The latter two had previously been determined using cytologic, biochemical, and genetic assays to form non-prion amyloid (Ngr1) or non-amyloid (Sla1) aggregates in yeast (Alberti et al., 2009; Sun et al., 2015). When analyzed by DAmFRET, all of the amyloid-forming proteins exhibited bimodality (Fig. S3A), although with dramatically different **δ** values that recapitulated their known prion-forming tendencies (Fig. 3A). For example, the nucleation barrier for Cyc8 PrD was more resistant to concentration increases than that of Mot3 PrD and Rnq1; the latter two proteins spontaneously form prions much more frequently than the former (Holmes et al., 2013; Liebman and Chernoff, 2012; Patel et al., 2009). Ngr1 PrL, the only amyloidogenic protein of the group that had not been found to form prions, had a smaller **δ** than all of the PrDs (Fig. 3A). Hence, the low prion propensity of Ngr1 PrL results not from an inability to form amyloids, but rather, an inability to ***not*** form amyloids when supersaturated. In other words, cells expressing it do not appreciably populate a “[*prion-*]” state. Upon close inspection, the data in Alberti et al. agree with this interpretation: Ngr1 PrL-Sup35C produced uniformly pink colonies at basal levels of ectopic expression, consistent with constitutive aggregation (Alberti et al., 2009). Finally, the Sla1 PrL acquired AmFRET in a predominantly concentration-dependent manner. That it lacked a detectable nucleation barrier suggests its assemblies are relatively disordered. Direct examination of the imaging data revealed that high AmFRET cells for all seven of these proteins contained puncta (Fig. S3B and data not shown).

**Figure 3.**
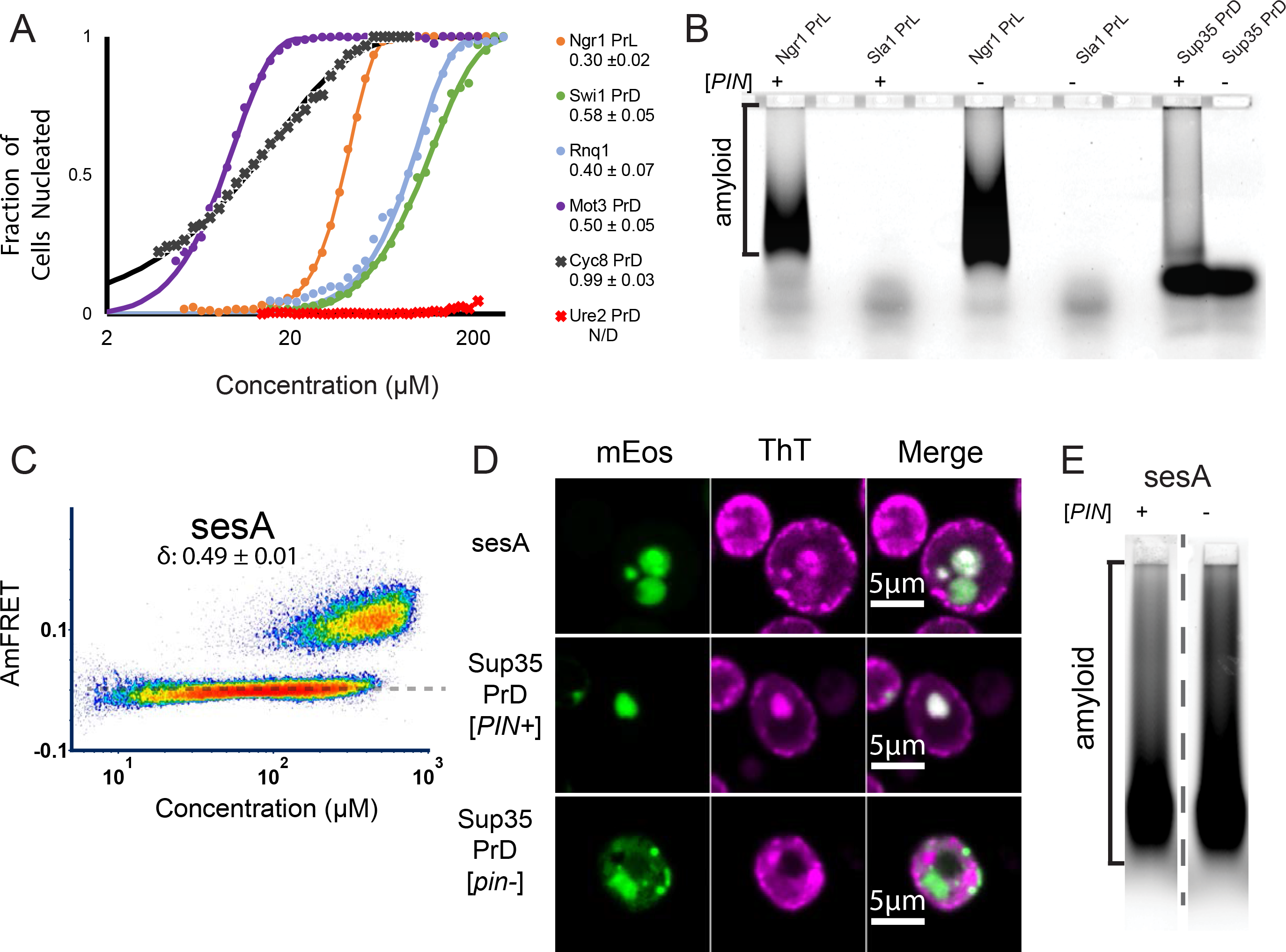
Nucleation barriers govern prion behavior. (A) Weibull fits of the DAmFRET of known amyloid-forming yeast polypeptides. The inset shows **δ** ± error. (B) SDD-AGE of Ngr1 PrL and Sla1 PrL in [*PIN+*] or [*pin-*] cells. (C) DAmFRET of sesA, showing an overlapping distribution characteristic of prions. (D) Representative images of ThT-stained cells expressing sesA, positive control Sup35 PrD in [*PIN+*] and negative control Sup35 PrD in [*pin-*]. ThT images were Gaussian-smoothed and adjusted for optimal contrast in ImageJ (see Methods). (E) SDD-AGE of sesA in [*PIN+*] and [*pin-*] cells. Images were background subtracted using a 250 pixel rolling ball and contrast adjusted for optimum band visualization (see Methods).

To directly evaluate the relationship of **δ** to the structural nature of the assembly, we examined the Ngr1 and Sla1 PrL puncta for hallmark features of amyloid. We stained live cells with thioflavin T (ThT), a molecular rotor that fluoresces upon binding repetitive features along the axes of amyloid fibrils (Biancalana and Koide, 2010; Kroschwald et al., 2015); and separately subjected them to Semi-Denaturing Detergent Agarose Gel Electrophoresis (SDD-AGE), a biochemical approach to detect polydisperse assemblies that resist denaturation in ionic detergents (Halfmann and Lindquist, 2008; Kryndushkin et al., 2003), a characteristic of amyloid that derives from its extensive hydrogen bonding network and exquisite side chain packing (Eisenberg and Sawaya, 2017). Cells expressing Ngr1 PrL contained ThT-positive foci and detergent-resistant polymers, whereas cells expressing Sla1 PrL contained ThT-negative foci and lacked detergent-resistant polymers (Fig. 3B, S3C).

Having successfully distinguished known prion from non-prion phase behavior, we next used DAmFRET for *de novo* prion discovery. The cytoplasmically inherited σ element of the filamentous fungus, *Nectria haematococca*, has been implicated by genetic and bioinformatic evidence as a prion of the sesA protein (Daskalov et al., 2012; Graziani et al., 2004). This prediction has not yet been tested. We therefore characterized sesA by DAmFRET. As a control for prion behavior by the same functional class of fungal prion proteins, we included the PrD of HET-s and its hypomorphic mutant W287A (Daskalov et al., 2014). Yeast cells expressing sesA or the WT HET-s PrD robustly partitioned into overlapping low and high AmFRET populations, exactly as shown for other prions (Fig. 3C, S3D). In contrast, much fewer cells expressing W287A HET-s PrD populated a high AmFRET state. Investigating further, we found that sesA formed ThT-positive puncta and detergent-resistant assemblies, as did the control protein, Sup35 PrD, when expressed in [*PIN+*] cells but not in [*pin-*] cells (Fig. 3B, D and E). In total, these data confirm an inherent capability of sesA to drive mutually exclusive cellular states through nucleation-limited amyloid formation, strongly suggesting that it is the protein determinant of σ.

### Nucleation barriers govern amyloid “cross-seeding”

Amyloids of one protein can increase amyloid formation by other proteins. Rnq1 prions, for example, enable the formation of Sup35 and Ure2 prions (Derkatch et al., 2001). The extent to which other prion proteins share this dependence, and its molecular underpinnings, remain unclear (Arslan et al., 2015; Keefer et al., 2017; Suzuki et al., 2012).

To clarify, we compared DAmFRET data for prion-like proteins in the presence ([*PIN*+]) and absence ([*pin-*]) of endogenous amyloid templates (Fig. S3A). Nucleation by Sup35 and Ure2 PrDs was undetectable in [*pin-*] cells, as expected. Swi1 PrD similarly depended on [*PIN*+], while Mot3 PrD exhibited a much less pronounced dependence, again as expected (Alberti et al., 2009). The ability to be templated by [*PIN*+] was unique to prion-forming proteins, as the two non-prion forming proteins nucleated independently of this factor, regardless of whether they formed amyloid (Ngr1 PrL) or non-amyloid (Sla1 PrL) assemblies. Cyc8 PrD also nucleated independently of [*PIN*+]. This, together with its exceptionally large **δ** (Fig. 3A), suggests that its self-assemblies differ structurally from those of other prions. Indeed, Cyc8 PrD features a highly unusual sequence element: a thirty-fold tandem repeat of the dipeptide motif, glutamine-alanine (Gemayel et al., 2015), which is expected to form self-assembling bilayers of amphipathic beta strands (Bowerman and Nilsson, 2012).

### A large nucleation barrier allows the archetypal prion protein, Sup35 PrD, to partition into physiological mRNP condensates

The apparent absolute dependence of Sup35 PrD amyloid nucleation on [*PIN*+], a cellular factor that does not exist in most cells (Halfmann et al., 2012), suggests that the amyloid state of Sup35 is unlikely to be physiological.

Many low complexity sequences have been found to undergo liquid-liquid phase separation that compartmentalizes protein activities in the cell (reviewed in Banani et al., 2017; Halfmann, 2016; Shin and Brangwynne, 2017). The molecular interactions in liquids are much weaker than those in amyloids; hence liquid-liquid phase separation generally only occurs at concentrations that are supersaturating with respect to amyloid formation (Halfmann, 2016; Vekilov, 2012). We reasoned therefore that Sup35’s endogenous existence at supersaturating concentrations (Tanaka et al., 2006) – and concomitantly its ability to form prions – implies a physiological propensity for liquid-liquid phase separation.

We noticed that [*pin-*] cells expressing Sup35 PrD to high concentrations exhibited slightly increased AmFRET versus cells expressing the fluorophore alone (Fig. 4A, S4A), which prompted us to take a closer look at the protein’s localization in the cells. We found that the AmFRET arose from small puncta in the cytosol. Some of these were larger than the resolution of our microscope (up to ~600 nm), allowing us to assess morphology (Fig. 4B-left). They were spherical (aspect ratio of 1.18 ± 0.02, n = 13). Amyloid puncta (in [*PIN+*] cells), in contrast, were highly aspherical (Fig. 4B-right).

**Figure 4.**
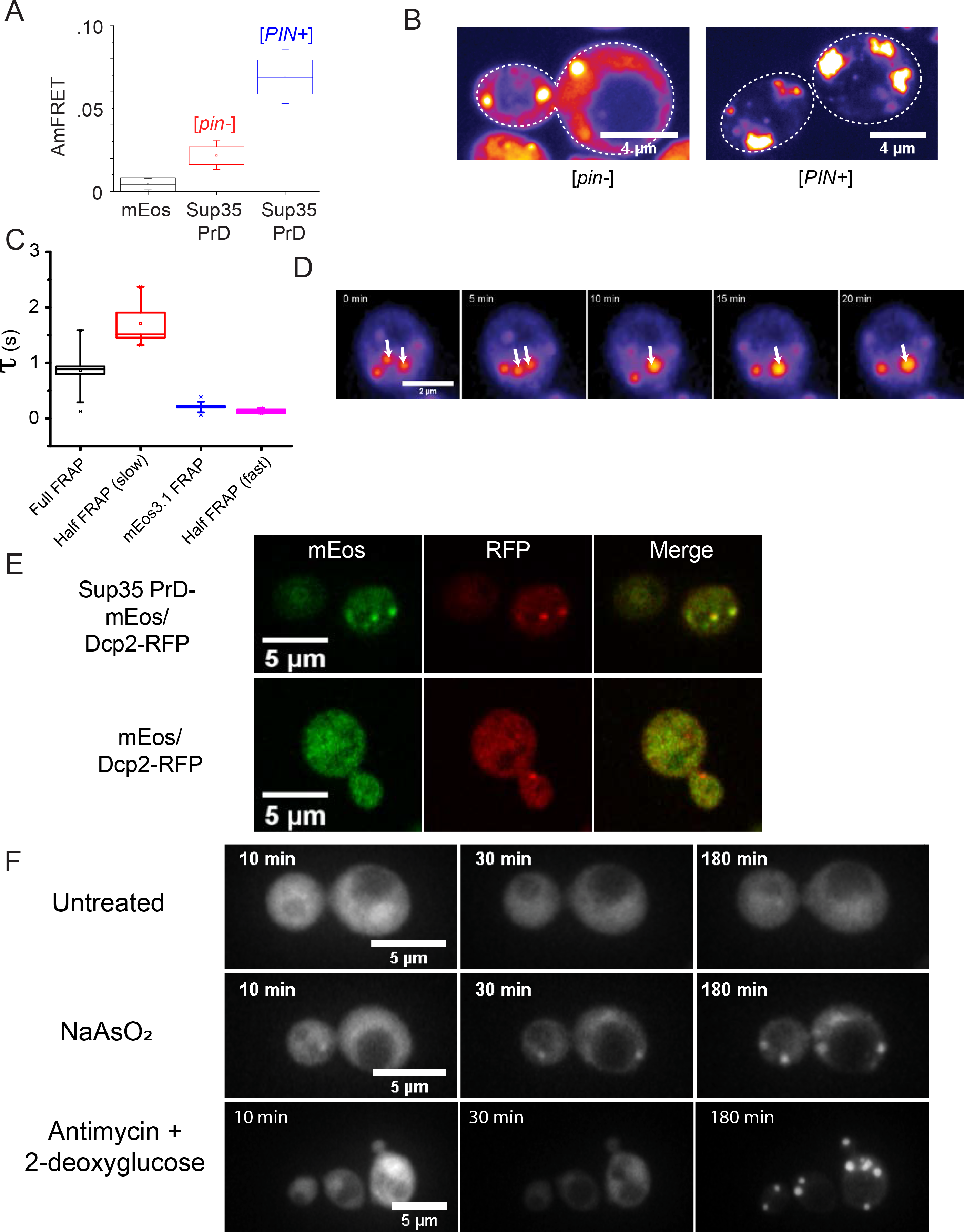
A large nucleation barrier allows the archetypal prion protein, Sup35 PrD, to partition into physiological mRNP condensates. (A) Box-Whisker plot showing that Sup35 PrD forms homotypic interactions even in [*pin-*] cells. The box is the SD of the mean, and the whiskers are the 5th and 95th percentiles of AmFRET values, for more than 2000 cells expressing between 80 and 200 μM of either unfused mEos3.1 (black) or Sup35 PrD in [*pin-*] (red) or [*PIN+*] cells (blue). (B) Representative confocal images of Sup35 PrD puncta in [*pin*-] (left) or [*PIN+*] cells (right). (C) Fluorescence recovery timescales of Sup35 PrD puncta in [*pin-*] cells as measured by FRAP. Error bars represent SD. (D) Time-lapse microscopy showing coalescence of Sup35 PrD puncta. Single confocal slice of a [*pin-*] cell expressing Sup35 PrD, showing two puncta (white arrows from 0-5 min) coalesce into one larger punctum (single arrow from 10 min). See also Fig. S4B. (E) Images of single confocal slices of cells co-expressing Sup35 PrD fused with mEos3.1 (and unfused mEos3.1 as control) and RFP-fused Dcp2 at 100X magnification. (F) Montage of cells expressing Sup35 PrD showing the formation of liquid droplets upon treatment with NaAsO_2_ (middle row) or antimycin + 2-deoxyglucose (bottom), versus untreated cells (top).

We used half-FRAP measurements to probe the internal dynamics of the spherical puncta. The half-FRAP recovery curves fit to a two component exponential, corresponding to recovery times of 0.275 ± 0.110 s and 1.710 ± 0.197 s for fast and slow components, comprising 9 ± 2 % and 28 ± 4 % of the recovery, respectively. The fast component approached that of cytosolic diffusion, 0.212 ± 0.008 s, as determined from cells expressing unfused mEos3.1. The slow component resembled that of whole punctum FRAP performed in the same cells, which yielded a recovery time of 0.864 ± 0.069 s. These data indicate that Sup35 PrD molecules diffuse almost as rapidly inside the puncta as they do in the bulk cytosol, but exchange relatively slowly across the interface between the two phases (Fig. 4C).

To further explore the nature of the Sup35 PrD puncta, we recorded their dynamics in living cells. The puncta were highly mobile. We observed two puncta within a cell coalescing into a larger punctum (Fig. 4D) whose fluorescence intensity exactly matched the sum of its precursor puncta (Fig. S4B). Note that the precursor puncta were significantly larger than the resolution of our microscope (diameters 410 nm and 500 nm). As a result, non-liquid contact between them would result in an elongated (aspherical) punctum. However, the coalesced punctum was instead spherical (aspect ratio 1.04, diameter = 490 nm). We conclude that Sup35 PrD partitions into liquid droplets when overexpressed in the absence of pre-existing Rnq1 and Sup35 amyloids.

Sup35’s PrD resembles that of LCSs that facilitate liquid-liquid phase separation by RNA binding proteins in the context of mRNP granules, such as P-bodies and stress granules. Full-length endogenous Sup35 localizes to these structures under heat stress (Grousl et al., 2013), and its PrD can substitute for the compositionally similar region of TIA-1 in targeting that protein to mammalian stress granules (Gilks et al., 2004). We therefore asked if Sup35 PrD droplets are, in fact, mRNP granules, by investigating their colocalization with a marker of both constitutive and stress-induced P-bodies: Dcp2 (Grousl et al., 2013; Rao and Parker, 2017). We observed that Sup35 PrD-mEos3.1, but not unfused mEos3.1, localized to Dcp2-RFP puncta (Fig. 4E). This finding rationalizes the low, concentration-dependent AmFRET of Sup35 PrD condensates – Sup35 PrD appears to be partitioning into endogenous condensates of other proteins, such that AmFRET becomes detectable only as it accumulates to high density within them.

To determine if Sup35 PrD phase behavior can be modulated by stress, we performed time-lapse microscopy on cells expressing Sup35 PrD-mEos3.1 exposed to 10 mM arsenite, a potent inducer of mRNP granules (Buchan et al., 2008). This treatment strongly promoted droplet formation (Fig. 4F, Movies S4-1, −2). To ensure that this observation was representative of a population of cells, we also quantified the number of cells positive for Sup35 PrD droplets before and after arsenite treatment for 1 hour, and found that 30% of the cells (22 out of 72) had droplets in the untreated, while 93% (69 out of 74) did upon arsenite treatment. We then asked if another mRNP granule-inducing stress – acute energy depletion via cotreatment with 10 μM antimycin and 20 mM 2-deoxyglucose (Riback et al., 2017) – produced the same effect. This condition, also, robustly induced Sup35 PrD droplets (Fig. 4F, Movies S4-3).

While this paper was in revision, Alberti and colleagues reported a tendency of modestly overexpressed Sup35 to form liquid droplets (Franzmann et al., 2018). To determine if endogenous Sup35 likewise partitions into non-amyloid assemblies under stress, we imaged chromosomally GFP-tagged full-length endogenous Sup35 by fluorescence microscopy. Neither confocal microscopy nor super-resolution microscopy was able to detect higher-order assembly of Sup35, even under conditions of acute energy depletion or arsenite stress (not shown). Given recent indications that a drop in cytosol pH may broadly underlie stress-induced phase separation (Munder et al., 2016; Riback et al., 2017), including that of full-length Sup35 (Franzmann et al., 2018), we then repeated the experiments under cytosol-acidifying conditions (see Methods). Still, no visible droplets formed. We reasoned that any assemblies that may exist must therefore be submicroscopic in size (<100 nm). To test for submicroscopic diffusing assemblies, we employed Fluorescence Correlation Spectroscopy (FCS). We found that when unstressed, Sup35 exists in two populations, one comprising rapidly- (*τ* _*D*_ ~ 0.5 ms) and the other slowly-diffusing (*τ* _*D*_ > 10 ms) species. Brightness analysis revealed that both populations contained an average of one Sup35-GFP molecule per diffusing particle (Fig. S4C). Given Sup35’s primary function in translation termination, these likely correspond to monomeric and ribosome-engaged molecules, respectively. When stressed by mild cytosol acidification, however, the average brightness of slow-diffusing particles increased two-fold, while the brightness of freely diffusing monomers remained unchanged (Fig. S4C). These data suggest that a fraction of Sup35 molecules redistribute into small multimeric species during stress. This interpretation agrees with recent reports that mRNP granules arise from the accumulation and coalescence of submicroscopic assemblies (Rao and Parker, 2017; Riback et al., 2017; Wheeler et al., 2016; Youn et al., 2018)

### Sup35 PrD mutants illuminate nucleation mechanisms in vivo

To explore potential linkages between liquid and amyloid self-assemblies, we compared phase behaviors with and without amyloid templates for a range of Sup35 mutants that have been identified in the literature to modulate Sup35 prion formation.

Two double mutants of the Sup35 PrD, Y46K+Q47K and Q61K+Q62K, had previously been found to be incompatible with certain amyloid isoforms (Bondarev et al., 2013, 2015). We observed that, in [*PIN+*] cells, these mutants greatly increased **δ** relative to WT (Fig. 5A, S5A, Table S1). This effect appeared to result, at least in part, from a reduced affinity of the proteins for themselves, as evidenced by a reduction in liquid-droplet associated AmFRET when expressed in the absence of Rnq1 amyloids.

**Figure 5.**
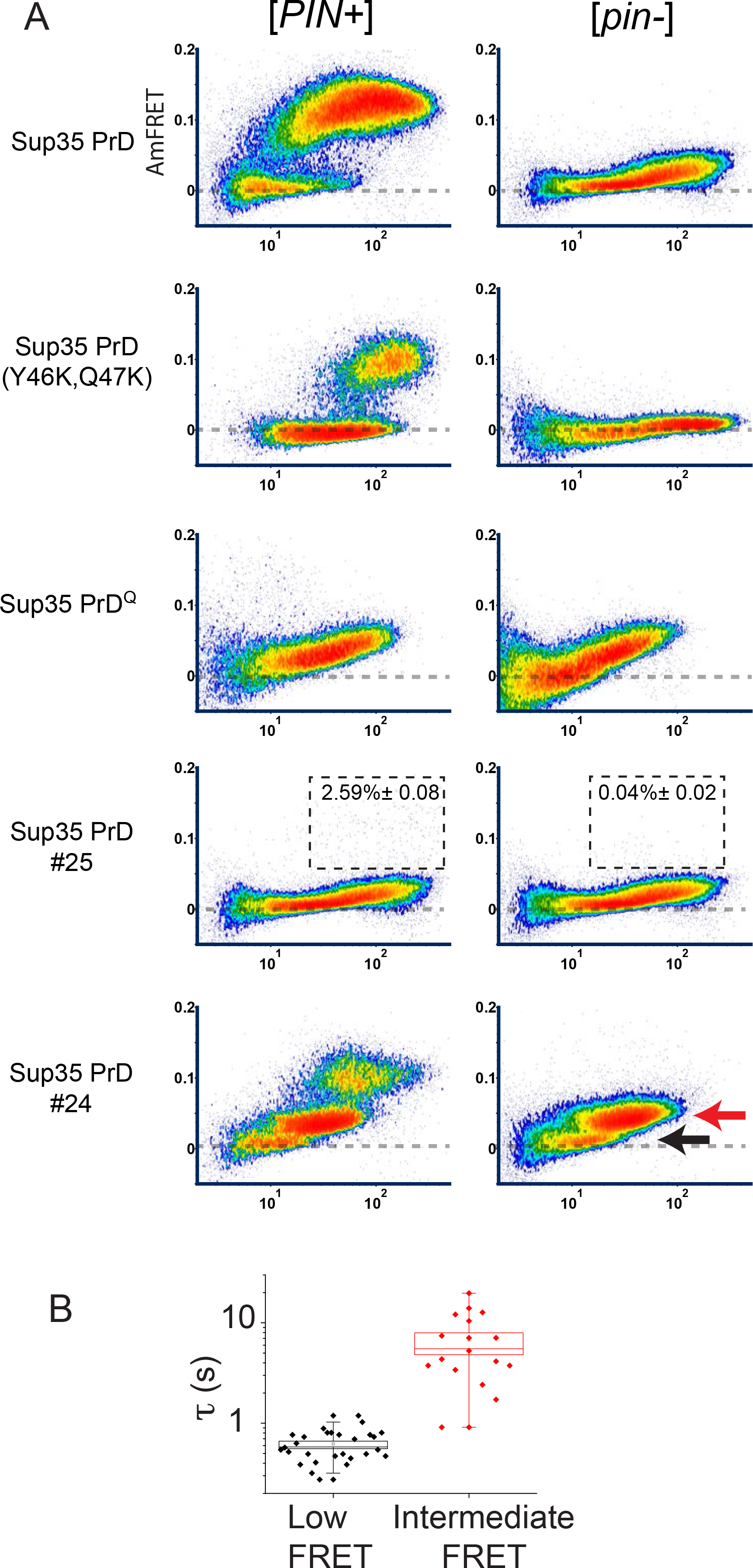
Sup35 PrD mutants illuminate nucleation mechanisms in vivo. (A) DAmFRET plots of select mutants and scrambled sequence variants (of overall identical amino acid composition) of Sup35 PrD in [*PIN+*] and [*pin-*] cells. Boxes in the plots for #25 designate the region considered prion-positive, with the percentage of total cells indicated. See also Fig. S5A. The red and black arrows indicate the “intermediate” and “low” AmFRET populations, respectively, that were sorted for microscopy analyses. (B) Quantification of fluorescence recovery times after half-puncta photobleaching for low (n = 28 cells) and intermediate AmFRET (n = 18 cells) states of [*pin-*] cells expressing Sup35 PrD #24. Boxes cover the SE, while the square inside the box shows the mean and the line shows the median. Whiskers delineate the 5th and 95th percentiles.

We next investigated more extreme changes in sequence. The Sup35 PrD is highly enriched for both glutamine (Q) and asparagine (N) residues. Despite their chemical similarity, the two side chains are not interchangeable with respect to amyloid formation. Replacing all of Sup35 PrD’s Qs with Ns (PrD^N^) increased prion propensity, while the reciprocal change (replacing all Ns with Qs; PrD^Q^) eliminated it (Halfmann et al., 2011). As expected, Sup35 PrD^N^ nucleated at lower concentrations than WT (below the sensitivity of DAmFRET; Fig. 5A, S5A).

The reciprocal mutant, PrD^Q^, produced an intermediate level of AmFRET that increased continuously with concentration (Fig. 5A). This distribution was not influenced by Rnq1 amyloids. Both features suggest non-nucleation-limited self-assembly, as shown above for Sla1 PrL (Fig. 3B, S3A, C). Indeed, ThT-staining and SDD-AGE confirmed an absence of amyloid-like ordered structure (Fig. S5C, D).

To determine if PrD^Q^ could nevertheless be templated to an amyloid state, we repeated DAmFRET in cells containing pre-existing amyloids of endogenous Sup35. Amyloids of a particular isoform ([*PSI+*^*strong*^], but not of [*PSI+*^*weak*^]; Tanaka et al., 2006) increased the AmFRET of PrD^Q^ (Fig. S5B), suggesting that the non-amyloid assemblies of PrD^Q^ are merely kinetically stable with respect to amyloid, and therefore represent “off-pathway” aggregates.

We next analyzed a series of Sup35 PrD variants that contain an identical amino acid composition as the WT protein, but with the order of residues scrambled (Ross et al., 2005). All of the scrambles were previously found to form prions. However, the prions of one of them (#25), were mitotically unstable, a phenotype previously linked to elevated thermodynamic stability (Tanaka et al., 2006). Given the dependence of the nucleation barrier on conformational ordering as demonstrated above, we predicted that #25 would exhibit a larger nucleation barrier than all of the other scrambled variants.

DAmFRET confirmed a very low nucleation frequency for #25 (approximately 2% of cells; Fig. 5A). Intriguingly, the protein’s acquisition of the low level of AmFRET attributable to condensation was unperturbed relative to WT, and persisted even in[*PIN+*] cells (Fig. 5A). This observation suggests that Rnq1 amyloids cross-seed other prion-like sequences through conformational templating, rather than increasing the affinity of the soluble proteins for themselves.

Two of the other scrambles, #21 and #26, populated low and high AmFRET states almost indistinguishably from WT, revealing comparable propensities for condensation and slightly decreased (#21) or increased (#26) propensities for amyloid nucleation (Fig. S5A).

The final analyzed scramble, #24, produced AmFRET distributions unlike any of the other Sup35 constructs. Cells that lacked pre-existing amyloid templates ([*pin-*]) partitioned into low and intermediate AmFRET populations (Fig. 5A). Cells that contained templates ([*PIN+*]) also partitioned into these populations, as well as a third population with a high AmFRET level indicative of the prion state.

We next used FACS to isolate low and intermediate AmFRET-containing cells, and characterized the state of #24 in each population using microscopy and FRAP. Rather than multiple small puncta with abundant diffuse fluorescence as we had observed for WT Sup35 PrD, both the low and intermediate AmFRET states of #24 corresponded to a single large punctum with virtually no diffuse fluorescence (Fig. S5C). The internal dynamics of the punctum differed dramatically between the two states: half punctum photobleached fluorescence recovered quickly in the low AmFRET state (albeit not as fast as for WT Sup35 PrD droplets), and very slowly in the intermediate AmFRET state (Fig. 5B). In neither population did the puncta stain robustly with ThT or contain detergent-resistant structure (Fig. S5C, D). Taken together these observations indicate that #24 partitions into a disordered condensate distinct from that of WT Sup35 PrD.

We speculate that the anomalous behavior of #24 could be attributed to an increased affinity of the protein for itself, resulting in larger, denser and more viscous puncta. The protein would be expected to have reduced miscibility with other components in the puncta (e.g. endogenous mRNP proteins). This could lead the protein to demix inside the puncta when expressed to very high levels, resulting in an even more viscous homogeneous phase (the intermediate AmFRET state). Such a multiphasic system has been documented within the liquid-like nucleoli of *Xenopus* oocytes (Feric et al., 2016).

To summarize, sequence changes to Sup35 PrD impacted amyloid nucleation in three distinct ways. 1) Y46K+Q47K and Q61K+Q62K reduced the affinity of soluble protein for itself, thereby reducing opportunities for productive conformational fluctuations. This is not surprising given the inhibitory effect of net charge on disordered protein self-solvation (Das and Pappu, 2013; Lin and Chan, 2017). 2) Consistent with lack of structure in dynamic condensates, scrambling the sequence of Sup35 PrD generally did not perturb the ability of soluble protein to interact with itself. It did, however, bias conformational fluctuations toward (#26) or away from (#21, #25) amyloid. 3) Finally, variants PrD^Q^ and #24 interfered with nucleation by routing the protein into “off-pathway” kinetically trapped assemblies.

### Nucleation barriers enable proteotoxic assemblies to accumulate

Despite their deep association with age-related degenerative disease, amyloids, themselves, are not overtly pathogenic. Rather, pathogenicity has been increasingly attributed to non-amyloid assemblies of the same proteins that are, fundamentally, metastable with respect to amyloid (Auer et al., 2012; Knowles et al., 2014; Serio et al., 2000; Vekilov, 2012; ten Wolde and Frenkel, 1997)

We noticed that Sup35 PrD^Q^ and #24 tended to express to lower concentrations than the other variants (Fig. 5A), suggesting they may be poorly tolerated by the cells. We had previously observed proteotoxicity of non-amyloid aggregates of PrD^Q^ (Halfmann et al., 2011). To test if #24 is likewise toxic, we plated serial dilutions of the cells to media that either induced or repressed ectopic protein expression. Variant #24, but none of the other scrambles, suppressed colony formation in [*pin-*] cells (Fig. 6A).

**Figure 6.**
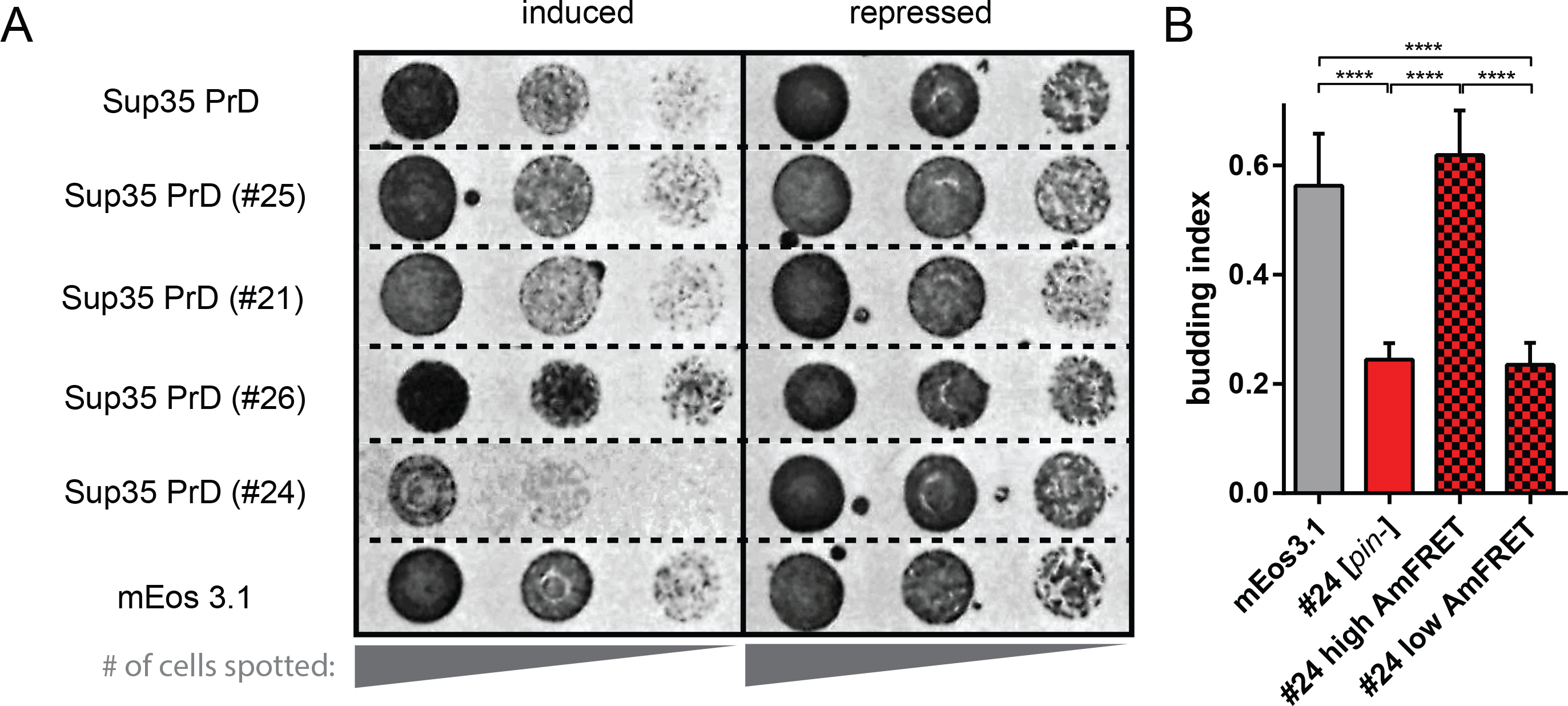
Nucleation barriers enable proteotoxic assemblies to accumulate. (A) [*pin-*] cells harboring plasmids encoding the indicated proteins were spotted as five-fold serial dilutions onto media that either induces (with galactose) or represses (with glucose) their expression. (B) Budding indices were calculated among cells expressing Sup35 PrD #24 or unfused mEos3.1 to the same intensity (5000 - 7000 AU). Solid red and checkered bars denote [*pin-*] and [*PIN*+] cells, respectively. The latter is divided into “high” and “low” AmFRET subpopulations corresponding to amyloid and non-amyloid assemblies, respectively. Shown are means of five experiments; error bars represent 95 % CI. ****, p<0.0001 (ANOVA).

To link toxicity to a specific state of the protein, we took advantage of the imaging capability of our flow cytometry setup to measure cell division, as a proxy for cell health, in populations of cells harboring different states of the protein. We accordingly repeated the DAmFRET experiment in the absence of cell cycle arrest, and used the brightfield channel in imaging flow cytometry to determine budding indices. Variant #24 severely reduced the budding index (Fig. 6B). Remarkably, the effect was entirely specific to its non-amyloid assemblies: the high AmFRET population of cells that occurred exclusively in the presence of [*PIN*+] budded just as frequently as cells expressing unfused mEos3.1. Hence, the amyloids ameliorated toxicity by draining the cell of kinetically trapped assemblies.

## DISCUSSION

We posited that exceptionally long timescales for diverse protein-driven phenomena result from kinetic barriers imposed by the nucleation of ordered self-assembly. We created a tool – DAmFRET – with the combination of features necessary to explore that hypothesis.

In short, DAmFRET employs a *single* fusion construct to produce a *direct positive* readout of protein self-assembly in each of thousands of *individual cells*. It has no restrictions on subcellular localization, solubility, or size of the assemblies, thereby mitigating false-negatives and false-positives. In contrast, existing high-throughput assays for protein aggregation report on large puncta (Narayanaswamy et al., 2009; Noree et al., 2010; Pereira et al., 2018; Ramdzan et al., 2012), inactivation of fusion partners (Alberti et al., 2009; Morell et al., 2011; Newby et al., 2017; Waldo et al., 1999; Zhao et al., 2016), require restrictive subcellular localization (Sivanathan and Hochschild, 2013), and/or necessitate expression from dual constructs (Arslan et al., 2015; Blakeley et al., 2012; Cabantous et al., 2013; Shyu and Hu, 2008). Most importantly, DAmFRET simultaneously reports total protein concentration, and thereby the concentration-dependence of self-assembly, in every single sample. This pivotal advantage allows for the detection and quantitation of nucleation barriers as bimodal dependencies of self-assembly on concentration.

Applying DAmFRET to proteins of structurally diverse self-assemblies, we discovered that many are indeed kinetically limited by nucleation barriers. These included all eight of the prion-forming domains tested, but not compositionally similar domains from non-prion proteins. Using bimodal DAmFRET as a diagnostic of prion behavior at the cellular level, we then validated a putative prion-forming protein that had been implicated genetically as the protein determinant behind a cytosolically-transmitted trait in a filamentous fungus. Moving forward, DAmFRET can complement measurements of cell-to-cell transmissibility *post*-nucleation, to provide a complete quantitative description of “prionness”.

### Deconstructing nucleation barriers

Our findings suggest that nucleation barriers relate closely to the structure of self-assemblies. Disordered assemblies lacked observable nucleation barriers, whereas ordered assemblies – amyloids, amyloid-like beta solenoids (HET-s PrD; Daskalov et al., 2014), and death fold filaments (ASC) – all exhibited nucleation barriers sufficient to enable deep supersaturation. As elaborated here, we believe this relationship follows directly from theory.

Nucleation barriers represent the combined improbability of fluctuations in each of the order parameters that distinguishes the new phase from the old. For protein self-assemblies, these can be summarized as density, orientation, and conformation. The probability of a critical density fluctuation depends directly on the concentration of the protein, whereas the other two parameters do not. Hence the more ordered a self-assembly is, relative to the unassembled phase, the more bimodal its DAmFRET. The value of **δ** therefore informs about the mechanism of nucleation, and hence, physical nature of the assembly.

Classical nucleation theory accounts for the contribution of the density fluctuation, which relates precisely to the degree of supersaturation as illustrated with phase diagrams. The nucleation barrier decays exponentially with supersaturation beyond the liquid-liquid coexistence line (green curves, Fig. 7; Vekilov, 2012).

Owing to their large sizes relative to the length scale of intermolecular interactions, protein liquid phases are believed to be inherently metastable with respect to crystallization (Lomakin et al., 1999, and references therein). Therefore, proteins within liquid droplets are theoretically supersaturated with respect to one or more ordered phases, whose boundaries lie at lower concentrations on the phase diagram (red curves; Fig. 7A-B). The nucleation barrier to ordered assembly is large between the two phase boundaries, where nucleation requires simultaneous fluctuations in both density and orientation (Asherie et al., 1996; Lomakin et al., 1999). The barrier may drop precipitously beyond the liquid-liquid coexistence line, however, where density fluctuations alone suffice to form condensates, within which the nucleation of ordered assembly becomes limited by an orientational fluctuation. As a result of this dependence, the phase diagram for crystallization of globular proteins can be approximated experimentally by determining the degree of supersaturation required for the induction of crystals under constant supersaturation rates (Asherie, 2004; Bhamidi et al., 2017; Nývlt, 1968; Peters, 2011). DAmFRET is a cytological analog of this approach.

**Figure 7.**
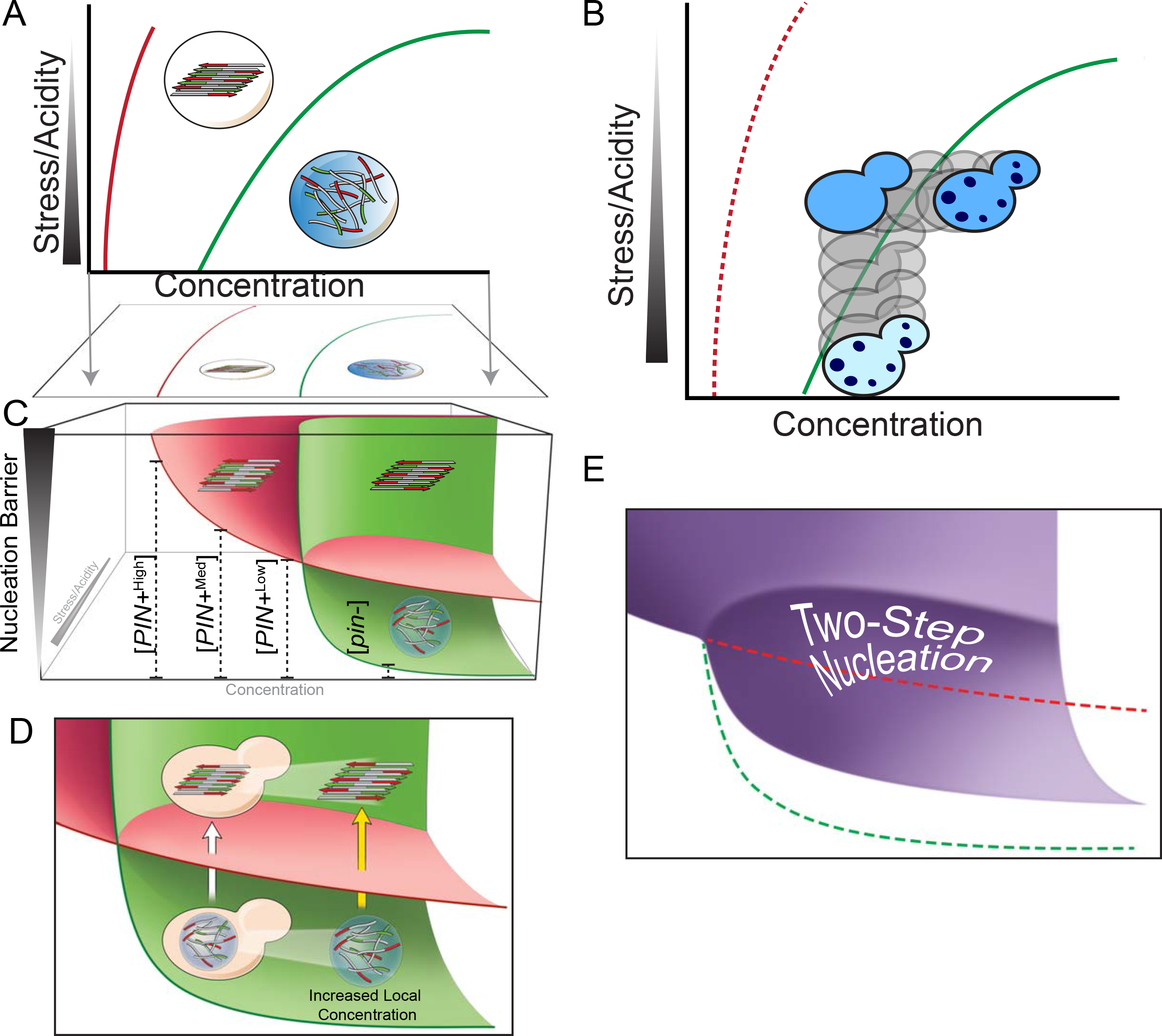
Nucleation barriers in phase diagrams. (A) Schematic protein phase diagram indicating the boundary thermodynamic conditions for ordered (e.g. amyloid) and disordered (condensate) self-assembly. Formally, each amyloid isoform will have its own phase boundary; for simplicity only one is shown (red). Amyloids tend to be more thermodynamically stable than disordered condensates; hence the phase space for amyloid encompasses that of condensation. (B) Schematic intracellular protein phase diagram for Sup35, with concentration on x-axis and stress/acidity on y-axis (increasing from top to bottom). The cell physiologically resides between the amyloid phase boundary (red dotted line) and liquid-liquid coexistence line (green line), and yet endogenous Sup35 remains dispersed over cellular timescales due to the large kinetic barrier to amyloid nucleation. Cells cross the liquid-liquid coexistence line to form condensates (green dots shown in cells) either upon exposure to stress (represented by travel downward) or upon experimental over-expression of Sup35 (horizontal travel). (C) Schematic of protein phase space that generalizes the findings of this paper. The phase diagram of (A) is projected along a third coordinate representing the magnitude of the kinetic barrier to nucleation. The red surface represents the barrier to amyloid nucleation, and the green surface represents the barrier to condensate nucleation. The surfaces intersect because amyloid nucleation depends less on density fluctuations than that of condensates (and hence declines less sharply with concentration). Heterogeneous templates (e.g. different [*PIN*+] isoforms), increase the probability of critical conformational changes, thereby allowing amyloid nucleation to be observed at lower concentrations. [*PIN*+] isoforms are arranged according to their observed reduction of Sup35 PrD **δ** (see also Fig. 2D and S2). (D) Illustration of two-step nucleation. The intersecting surfaces illustrate that, at concentrations above the liquid-liquid coexistence line, amyloid nucleation will tend to proceed through condensed intermediates. Condensation of the protein into droplets increases its local concentration, as indicated by the rightward shift of the droplet, reducing its barrier to amyloid nucleation as indicated by its closer proximity to the red surface. The white arrow depicts the transition to amyloid at the average concentration of the protein in the cell, while the yellow arrow depicts the transition occurring at the effective concentration resulting from condensation. (E) The theoretical observed dependence of the amyloid nucleation barrier on concentration resulting from two-step nucleation as described in (D).

Amyloids further differ from globular protein crystals in that their nucleation requires fluctuations not only in density and orientation, but in *conformation* as well. Hence a rate-limiting barrier to amyloid nucleation may persist even at concentrations well above the liquid-liquid coexistence line (“two-step nucleation”; Fig. 7D-E). Indeed, amyloid nucleation *in vitro* is preceded near universally by an accumulation of metastable “oligomers” (Auer et al., 2012; Chatani and Yamamoto, 2017; Serio et al., 2000). These can be either on-pathway, as described above for crystals, or off-pathway, if their internal viscosity sufficiently reduces conformational fluctuations or if they refold into local energetic minima (Vitalis and Pappu, 2011).

Our data suggest that conformational fluctuations also broadly dominate the amyloid nucleation barrier *in vivo*. Sup35 PrD, a model amyloid-forming protein, formed visible droplets that – based on fluorescence intensity and size – contain millimolar concentrations of monomer. This corresponds to at least ten thousand-fold supersaturation with respect to the concentration of monomer that remains dispersed in amyloid-containing cells (approximately 50 nM; Tanaka et al., 2006). And yet, even though it remained fully mobile within the droplets, the protein failed to acquire the thermodynamically favored amyloid state over experimental timescales.

Instead, our data showed that amyloid nucleation by Sup35 as well as Ure2 and Swi1 PrDs depended virtually entirely on the existence of pre-formed amyloids of another protein. Those heterologous amyloids did not change the affinity of soluble prion-forming proteins for themselves, as they did not increase partitioning into pre-amyloid condensates (Fig. 5A and S3A). Therefore, cross-seeding occurred by specifically increasing the probability of critical conformational changes, without affecting other components of the nucleation barrier, which corresponds a large, medium, and small **δ** for Sup35 PrD in the presence of low, medium, and high [*PIN*+], respectively (Fig. 7C). This interpretation rationalizes the extraordinary sequence-dependence we observed for cross-seeding by [*PIN+*], which differed dramatically even among proteins with overtly similar amino acid compositions (Figs. 3, S3, 5 and S5).

Altogether, our data indicate that intracellular nucleation of disordered condensates occurs through a critical fluctuation in polypeptide density that occurs too frequently to restrain self-assembly kinetically. Nucleation of ordered polymers, in contrast, occurs via fluctuations in polypeptide density, orientation and conformation. The simultaneous occurrence of critical fluctuations in all these parameters appears to be sufficiently improbable as to prevent nucleation over cellular timescales. We believe this is why all well-characterized prions manifest as highly ordered polymers. The improbability of acquiring that level of order, from a disordered dispersed state, produces a nucleation barrier large enough for prion-free cells to harbor fully soluble protein despite being supersaturated at endogenous concentrations.

### Protein over-expression unmasks latent phase behavior

That disordered condensation lacks an appreciable nucleation barrier allows it to respond rapidly to small changes in thermodynamic parameters such as temperature, pH, and ligand concentrations. Does nature exploit this? The astonishing variety of cellular processes now known to be modulated by liquid-liquid phase separation indicates that indeed it does, and to great extent (Banani et al., 2017; Falahati and Wieschaus, 2017; Franzmann et al., 2018; Riback et al., 2017)

Phase boundaries can be crossed by moving along either axis of a phase diagram. We reasoned that by pushing protein concentrations well above endogenous levels, we can systematically cross phase boundaries that may, nevertheless, be physiological in other dimensions, such as pH (Fig. 7A). DAmFRET validated this idea for Sup35 PrD. The protein condensed into non-amyloid assemblies not only at high concentration during stress-free growth, but at lower concentrations under stress, which induces mRNP granules through cytosol acidification. These colocalized with mRNP granules, suggesting that Sup35 partitions alongside other mRNA-associated proteins into physiological condensates of heterogeneous composition (De Leeuw et al., 2007; Gilks et al., 2004; Tourrière et al., 2003; Wilczynska et al., 2005). Alberti and colleagues largely corroborate these findings (Franzmann et al., 2018): Sup35-GFP, when modestly overexpressed from a low copy number plasmid (Karim et al., 2013), formed liquid droplets during stress. These protected the essential enzymatic activity of Sup35 from inactivation by those stresses.

Nevertheless, over-expressing a protein does more than change its absolute concentration in the cell. It also changes its concentration *relative* to binding partners, and may induce cellular responses; both effects can influence phase behavior. We therefore went a step further, and verified that endogenous Sup35 tagged with GFP also forms multimers during stress.

“Prion-like” describes sequences that resemble the canonical yeast prion proteins: Ure2, Sup35, and Rnq1 (Alberti et al., 2009; Cascarina and Ross, 2014). They tend to be lengthy and replete with polar uncharged residues. These features do not directly promote amyloid (Goldschmidt et al., 2010; Knowles et al., 2014; Maurer-Stroh et al., 2010). Instead, they promote dynamic condensation by increasing low affinity polyvalency (Banani et al., 2017; Halfmann, 2016). Indeed, most prion-like sequences exhibit no detectable propensity to form amyloid (Alberti et al., 2009), and we found that randomly scrambling Sup35’s PrD reduced its fluidity – either directly (#24) or by facilitating amyloid nucleation (#26). We strongly suspect that prion-like sequences evolve not to promote aggregation thermodynamically, but rather, to prevent aggregation kinetically. The consequence, at least for several such proteins, appears to be catastrophic self-sustaining aggregation once the kinetic barrier is breached.

### Condensate metastability underlies proteopathic aggregation

The single cell resolution of DAmFRET enabled us to query the relationship of phase behavior to proteotoxicity. Specifically, we found that a variant of Sup35 PrD (#24) destabilized the supersaturated soluble state by allowing the protein to form viscous non-amyloid condensates. Remarkably, cells containing these condensates suffered a growth defect; and lowering the nucleation barrier to amyloid mitigated toxicity by draining the protein from those assemblies into the benign and more thermodynamically favored amyloid phase. The extent to which human amyloid-associated proteopathies likewise result from kinetically trapped non-amyloid assemblies remains to be tested. Nevertheless, our finding supports an emerging paradigm that non-amyloid assemblies contribute more to pathology than do amyloids (Halfmann, 2016; Knowles et al., 2014)

### Concluding remarks

Unlike protein folding at the secondary, tertiary, and quaternary levels, phase transitions can be rate-limited over biological timescales by nucleation. The larger the nucleation barrier, the more discrete the protein’s activity in space and time. Nucleation-limited self-assembly therefore enables switch-like physiological changes that are simply not possible with proteins acting individually. Nature has acted upon protein kinetics at the ensemble level.

Some features of self-assembly structure may have therefore evolved according to its effects on nucleation barriers, rather than enzymatic activities and ligand interfaces that tend to shape protein evolution at lower structural levels. This implies relationships between structure and function that may otherwise seem superfluous. Disordered protein condensates tend to have small nucleation barriers that limit their functions to subcellular spatiotemporal scales. In contrast, ordered assemblies such as amyloids have large nucleation barriers enabling them to function over organismal and population-level spatiotemporal scales. To some extent, then, the material existence of a particular assembled structure may be irrelevant to its function. Testing this idea, its generality among nucleated self-assemblies, and its potential contribution to their time-dependent cellular activities, presents a fantastic challenge for the future. The methods developed here provide ways to overcome this challenge.

## Author Contributions

Conceptualization, T.K., T.S.K., E.K., S.V., and R.H.; Methodology, T.K., T.S.K., E.K., S.V., and R.H.; Investigation, T.K., T.S.K., J.W., E.K., S.V., J.J.L., A.R.G.; Formal Analysis, T.K., T.S.K., J.W., E.K., S.V., J.J.L., A.B., J.R.U., and R.H.; Data Curation, M.C.; Visualization, T.S.K., J.J.L., and R.H., Writing – Original Draft, R.H.; Writing – Review & Editing, M.C., S.V., T.S.K., J.W., J.J.L., J.R.U. and R.H.; and Funding Acquisition, R.H.

## Acknowledgements

We thank Susan Liebman and Jonathan Weissman for variant [*PIN*+] and [*PSI*+] yeast strains, respectively; Eric Ross for scrambled Sup35 PrD cloning templates; Roy Parker for plasmid pRP1156; and Ammon Posey, Rohit Pappu, and Boris Rubinstein for insightful discussions. We thank Mark Miller and Chris Wood for assistance preparing illustrations, and Michelle Tan for assistance with molecular biology. This work was funded by NIH Director’s Early Independence Award DP5-OD009152, NIH grant P30 AG035982, and the Stowers Institute for Medical Research.

## Methods

### Cloning procedures

A gateway destination vector, BB5b, was constructed by ligating a GeneArt String encoding a yeast codon-optimized 4x(EAAAR) linker and mEos3.1 between HindIII and XhoI in pAG426GAL-ccdB (14155). The *URA3* promoter was truncated to increase plasmid copy number (Loison et al., 1989). A golden gate (Engler and Marillonnet, 2013) cloning-compatible vector, V08 was constructed from BB5b using gap repair to replace the Gateway cassette with inverted BsaI sites. V08 was then used to construct V12 by ligating a synthetic fragment encoding yeast codon-optimized mEos3.1-4x(EAAAR) followed by inverted BsaI sites between SpeI and XhoI. Finally, vector CA was constructed from V12 using gap repair to replace the inverted BsaI sites with the Gateway cassette.

Inserts available as pre-existing Gateway entry clones (Alberti et al., 2009; Cai et al., 2014; Douglas et al., 2008; Halfmann et al., 2011) were introduced into BB5b and CA using Gateway LR recombination. All other inserts were ordered as GeneArt Strings flanked by Type IIs restriction sites for ligation between self-excising BsaI sites in V08 and V12. All plasmids were verified by sequencing. Table S2 lists the plasmids and encoded polypeptide sequences for all fusion proteins characterized in this study.

### Yeast genetic manipulations

Table S3 details all yeast strains used in this study. Yeast were transformed with a standard lithium-acetate protocol (Gietz et al., 1992). The primary DAmFRET strain, y1713, was constructed from Y7092 (Tong and Boone, 2007). PCR-based mutagenesis (Goldstein and McCusker, 1999) was used to replace *CLN3* in its entirety with a purpose-built cassette that expresses *WHI5* from the inducible *GAL1* promoter. Strains y1851 and y1852 were constructed by passaging strains Y7092 and y1713, respectively, four times on YPD plates containing 3 mM GdHCl, a prion-curing agent (Ferreira et al., 2001). Toxicity and budding index analyses were performed in Y7092 and y1851.

*GAL1* promoter-mediated overexpression of *WHI5* in a *cln3*-knockout background potently arrests cells in G1 phase (Adames et al., 2015), thereby preventing nucleated protein assemblies from transmitting beyond the original cell, while enabling more accurate calculation of cell volume due to the spherical shape of the arrested cells. Growing the yeast in glucose-based medium enables the cells to proliferate, whereas switching them to galactose-medium induces arrest and simultaneous induction of fusion protein expression.

### Preparing cells for DAmFRET

Standard yeast media and growth conditions were used. Single transformant colonies were inoculated to 200 µl of glucose-containing selection medium per well in a round bottom microplate, then incubated on a Heidolph Titramax vibrating platform shaker at 30℃, 1350 RPM overnight, to allow for the prevalence of a range of copy numbers of plasmid in the population and to obtain a turbid culture. Cells were then washed twice with sterile distilled water to remove residual glucose before being resuspended in 200 µl of galactose-containing induction medium and returned to the incubating shaker for approximately 16 hrs. Microplates were then illuminated with an OmniCure^®^ S1000 fitted with a 320-500 nm (violet) filter and a beam collimator (Exfo), positioned 45 cm above the plate, for a duration of 25 min, which was found to produce the maximum acceptor fluorescence with minimal photobleaching of donor. Violet light induces cleavage in the mEos3.1 peptide backbone adjacent to the chromophore (Wiedenmann et al., 2004; Zhang et al., 2012), converting it from a green form (emission peak at 516 nm) to a red form (emission peak at 581 nm). The beam power at the plate was 11.25 mW/cm^2^, giving a total photon dose of ~17000 mJ/cm^2^. Microplates were shaken at 800 RPM on a microplate shaker during photoconversion to prevent cell settling.

### DAmFRET Data Collection

All the AmFRET data were acquired on an ImageStream^®x^ MkII imaging cytometer (Amnis) at 60X magnification with low flow rate and high sensitivity using INSPIRE software. INSPIRE software directed the instrument to acquire as follows: channel 04 (brightfield in camera one), channel 10 (brightfield in camera two), channel 02 (donor fluorescence), channel 03 (sensitized emission FRET), channel 07 (blue fluorescence, a proxy for dead/dying cells as validated by staining with Sytox Far Red; see Fig. Supplemental 1A) and channel 09 (acceptor fluorescence). Magnification at 60X provided a pixel size of 0.3 μm^2^. All samples were loaded from the microtiter plate using the ImageStream^®x^ MkII autosampler. Channels 02 and 03 captured emission from 488 nm excitation, with 528/65 and 577/35 nm filters, respectively. Channel 07 captured emission from 405 nm excitation, with a 457/45 nm filter. Channel 09 captured emission from 561 nm excitation, using a 582/25 nm filter.

Brightfield-based gates were assigned in INSPIRE: first for focused events, determined by gradient root mean squared, and second for single cells, determined by area and aspect ratio (ratio of the lengths of the long and short axes through the cell). These focused, single-cell events were counted for donor and acceptor fluorescence positivity. For each sample, a minimum of 20,000 double positive events or maximum of between 5 and 10 minutes collection time were counted before proceeding to the next sample. Although only putative target events were counted, all unsaturated events were acquired and saved.

Compensation of the data collected was performed by using the built in wizard of IDEAS 6.2 on single color controls – cells expressing non-photoconverted mEos3.1 and those with dsRed2 (as a proxy for the red form of mEos3.1).

### DAmFRET Data Analysis

Data were processed using IDEAS 6.2 (Amnis) software and batched using FCS Express Plus 6.04.0015 software (De Novo). IDEAS yields standard parameters, such as integrated intensity of acquired channels, as well as user-derived features, such as AmFRET (FRET intensity/acceptor intensity). To measure cell area from brightfield, we created a feature for area calculated by the adaptive erode mask, with an adaptive erosion coefficient set at a threshold of 70%, which both visually aligns with the cell boundary and corresponded to mean cell area from a culture simultaneously measured by microscopy. All integrated intensity values reported or intensity derived features (e.g. cytosolic concentration) exclusively represent intensity within this brightfield mask.

AmFRET positive population fractions were determined by dividing cytometry histograms into 64 bins logarithmically spaced from 1 to 1000 micromolar. For each protein, the threshold for the AmFRET positive population was measured as the point halfway between the two population centers as determined by a multi-Gaussian fit of the AmFRET distribution. For strains where only the positive or negative population are observed, the threshold value was determined by a closely related protein showing both populations. For each bin, the fraction of cells in the AmFRET positive population was determined. Bins at the low and high extremes of concentration were excluded when their fractions deviated above and below, respectively, neighboring bins due to low event number and autofluorescence. These curves were then fit to a Weibull function of the following form, using non-linear least squares optimization via the Levenberg-Marquardt method (Bevington and Robinson, 2003):,

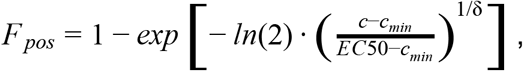

where *c* is the concentration and *EC50* is the concentration at which the AmFRET positive population is equal to 50%. The shape parameter or “Weibull slope”, *1/***δ**, describes the sharpness of the transition. We therefore use its reciprocal (**δ**) to report the observed persistence of the nucleation barrier with respect to concentration. A two parameter Weibull function was previously used to phenomenologically describe nucleation as a function of supersaturation (Sear, 2016). Since we do not know the saturating concentrations of each self-assembly *a priori*, we included a third parameter, *c*_*min*_, to allow *F*_*pos*_ to go to 0 as required for phase behavior as a function of raw concentration rather than degree of supersaturation. Errors were obtained by the monte carlo method, employing random noise added to the best fit curve with a standard deviation equal to the fit residuals standard deviation. 100 such randomized curves were fit to obtain standard errors in the fit parameters.

DAmFRET is highly analogous to isothermal metastable zone width (MZW) measurements used to describe the range of supersaturating concentrations under which crystallization can be induced. MZWs are determined empirically and the precise value depends on the rate of supersaturation (Asherie, 2004; Bhamidi et al., 2017). They can be used to approximate phase diagrams provided crystal nucleation occurs rapidly within disordered clusters (Asherie et al., 1996). Our cellular system does not offer the level of experimental control (for example, constant supersaturation rates in all vessels), or the level of precision in determining saturating concentrations, that would be required to confidently link DAmFRET with MZWs. Moreover, because amyloid nucleation (in contrast to crystallization of well-behaved globular proteins) can be rate-limited by conformational fluctuations even within condensates, the metastable zone (usually defined as the region between the liquidus and binodal) will have limited relevance to amyloid nucleation barriers (as elaborated in Discussion and Fig. 7).

### Determining absolute protein concentration from fluorescence intensity

Molecular brightness of photoconverted mEos 3.1 was calibrated by ImageStream ^®x^ MkII measurement of mEos3.1 endogenously fused to Spc42, a protein in S. cerevisiae with about 1000 assembled molecules per cell (Bullitt et al., 1997), in order to relate instrumental intensity values to molecule number. Our calibration method is based on methods from fluorescence correlation spectroscopy (Müller, 2004; Shivaraju et al., 2012; Slaughter et al., 2011). Brightfield area measurements generated by the instrument permitted extrapolation from cell size and Spc42-derived molecular brightness to gross cellular concentration. Given that organelles occupy about 17% of haploid S. cerevisiae cell volume (Uchida et al., 2011), we corrected the calculation to reflect concentration from fluorescence intensity generated in the cytosol.

The method firstly consists of the measurement of unconverted mEos3.1 molecular brightness as defined by the peak intensity observed by a single mEos3.1 molecule using Spc42 as a reference standard. One attractive feature of the spindle pole body is that its size of ~150 nm (Bullitt et al., 1997) is significantly smaller than the resolution of the imaging cytometer. As a result, the observable spot on the imaging cytometer is the same size as would be expected from a single fluorescent molecule and its peak amplitude will be proportional to a single molecule brightness multiplied by the fluorophore copy number.

Cells expressing Spc42-mEos3.1 were acquired on the ImageStream at 40, 80, 160, and 400 mW 488 nm laser powers. The IDEAS software was utilized to filter for unbudded and live cells based on scatter plots of area vs. aspect ratio and fluorescence intensities in ch02 vs. ch07, respectively. Compensated images were exported as 16-bit tiff images for visualization and further analysis with a custom implementation of the Bio-Formats plugin for ImageJ. Spc42 spots were clearly resolved in approximately 10% of the cells. The cells without clearly resolvable spots could have been out of focus. The maximum intensity in each image was selected as the initial center point of the Gaussian function. Gaussian fitting for the maximum spot in each image was then accomplished with a custom grid search fitting algorithm in ImageJ available at http://research.stowers.org/imagejplugins. The plugin attempts to fit a Gaussian function centered at each 0.25 pixel increment within 2 pixels in either direction from the maximum and with standard deviations from 0.5 pixels to 4 pixels at 0.1 pixel increments. At each candidate grid point, the fit is performed with linear least squares to a Gaussian function as follows:,

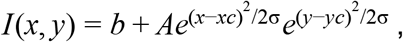

where *xc* and *yc* are the candidate centers of the actual spot, *σ* is the candidate spatial standard deviation. Linear least squares solves for the background intensity, *b*, and the peak amplitude, *A* at each candidate grid point. The point with the lowest *χ*^2^ value is chosen as the best fit.

After fitting, a 2D histogram of peak amplitude vs. standard deviation reveals two behaviors. At small standard deviations there is a downward trending shoulder showing decreasing amplitude with increasing standard deviation. This is expected behavior for particles at different focal planes of the microscope. At higher standard deviations, the amplitude does not depend on standard deviation and these spot sizes correspond to values much larger than the expected resolution of the image cytometer (> 1 µm). This seems to correspond to the ~90% of cells that did not show visible spots as mentioned above.

A histogram was made of peak amplitudes with standard deviations less than 1.1 pixels (widths less than 850 nm). We estimate the standard deviation of the smallest spots to be 0.9 pixels (width of 700 nm), so this allows for a narrow distribution of spot sizes around the minimum value. These histograms as a function of laser power are shown in Fig. S1E. This histogram was then fit to a one dimensional gaussian function, this time using traditional non-linear least squares to obtain the center of the peak amplitude distribution. At 40 mW, the Spc42 peaks become more difficult to resolve and the peak amplitude distribution appears bimodal. The higher amplitude peak follows the trend from the other laser powers and represents “real” Spc42 spots.

The relationship between Spc42 peak amplitude and laser power is not linear (Fig. S1E). In order to create a calibration curve for different laser powers, the center of the peak amplitude distribution as a function of laser power was fit to a simple exponential function as follows:,

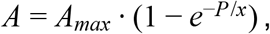

where *P* is the laser power, *A* is the peak amplitude, and *A*_*ma*x_ and *x* are fit variables. For our setup, *A*_*max*_ and *x* are 70.16 and 226.1, respectively. The brightness per mEos3.1 molecule is then obtained by simply dividing the *A*_*max*_ value by 1000 molecules. Our brightness per molecule for unconverted mEos3.1 is therefore 0.00594.

Now that we can derive the molecular brightness of green unconverted mEos3.1 at each laser power, we can use the integrated intensity from each cell to calculate the fluorophore concentration. Firstly, we use the measured area of each cell (from the Ideas software) in pixels to calculate the average intensity per pixel. From fluorescence correlation spectroscopy theory, we know that the average intensity is the product of the number of molecules in the focal volume multiplied by the molecular brightness of each molecule. In turn, the number of molecules per focal volume can be converted to concentration as follows:,

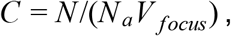

Where *N*_*a*_ is Avogadro’s number and *V*_focus_ is the microscope focal volume. The latter can be somewhat difficult to determine because of the poorly defined z dimension of the imaging cytometer focus. If we measure the average peak intensity of sub-resolution beads at differing shifts of the objective from the center of the flow stream, we obtain an approximate focal volume profile (Fig. S1D) with a width of 4.2 µm. Therefore we have chosen to treat the focal volume as a cylinder with radial gaussian profile of width 700 nm and a z extent of 4.2 µm, giving a focal volume of 2.7 µm ^3^ or 2.7 fL.

The final piece is to calibrate the photoconversion efficiency. We have chosen to express our concentrations in unconverted mEos3.1 concentration units as measured in the green channel. As a result, we can measure the average ratio of the intensity of a photoconverted cell in the red channel to an unconverted cell in the green channel. That ratio allows us to convert photoconverted red intensity into unphotoconverted green intensity for every sample and subsequent concentration calibration. In addition, we assume that most of the proteins analyzed will be cytoplasmic. The ratio of cytoplasmic to total volume in a yeast cell has been estimated as 0.83 (Uchida et al., 2011).

To summarize, cytosolic concentration (*C_cytosol_*) assessment contains three derivations.

1. Determine the relationship between laser power and photoconverted donor intensity of a 1000 molecule complex
2. Find the conversion factor from integrated intensity to concentration given optic constraints (especially focal volume) of the instrument’s 60x objective
3. Relate the photoconversion ratio of photoconverted green intensity to photoconverted red intensity of the mEos-only control

This last piece is crucial given that AmFRET signal accompanying mEos association is necessarily accompanied by loss of donor signal, whereas acceptor is constant and thus is an appropriate, AmFRET signal independent measure of fluorescent mEos. We can simplify the derivation above to represent the average concentration in the cytoplasm of a cell as:

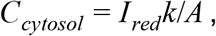

where *I*_*red*_ is the compensated cell intensity in the red channel and *A* is the cell area in µm^2^. Here, the multiplier, k, is as follows for 20 mW laser power:

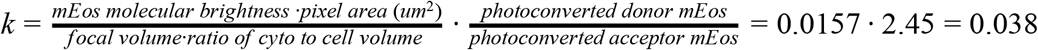

### Molecular Brightness Analysis

Fluorescence correlation spectroscopy was used to determine the molecular brightness and thus the extent of oligomerization of Sup35-EGFP under different conditions (Fig. S4B). Brightness analysis was performed in an analogous fashion to (Slaughter et al., 2013). Briefly, the amplitude of an FCS curve can be written in terms of the number of molecules per focal volume, *N*, and the molecular brightness, *ε*:

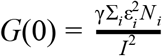

Here *ɣ* is a shape factor accounting for the shape of the confocal focal volume. It’s value is approximately 0.27 for most confocal microscopes (Slaughter et al., 2013) but given our desire for relative molecular brightness, it doesn’t matter for this study. The average fluorescence intensity can be expressed as follows:

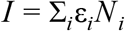

The measurement of the total FCS amplitude and average intensity doesn’t provide enough information to measure more that the average brightness and number of molecules. However, the presence of multiple diffusion times affords the opportunity to measure two amplitude values:

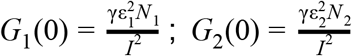

Substituting in the previous equation, we can express this in terms of the fractional intensity, *f*, of the fast diffusing component:

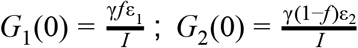

Rearranging we can get a “brightness” curve with amplitudes as follows:

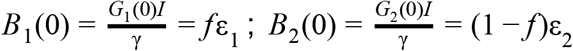

This leaves us with three unknown components. If we assume that the fast component is monomeric (brightness equal to GFP monomer), we can easily solve for the fractional intensity and the slower component brightness.

Error bars in correlation amplitudes represent standard errors in the fitted parameters and were obtained by monte carlo analysis as with the DAmFRET population analysis (see above). Errors were propagated to brightness values using standard derivative formulas ignoring covariances between terms (Bevington and Robinson, 2003).

### Confocal imaging, time-lapse, and FRAP Analysis

Confocal images were acquired on an Ultraview Vox Spinning Disc (Perkin Elmer). The green form of mEos3.1 was excited using a 488 nm laser through either an alpha Plan Apochromat 100x 1.46NA objective (Zeiss) or a Plan Apochromat 63x 1.4NA objective (Zeiss). Emission was collected with an EMCCD (Hamamatsu, C1900-23B) and was filtered with a 525-50 nm bandpass filter. Movies were acquired with cells trapped in a CellASIC Onix2 microfluidic device (Millipore) with either single confocal slices or z-stacks. The z step spacing for movies was set to between 0.3 and 1 µm. The time scale between images varied. FRAP was acquired on the same system in FRAP preview mode, with 5 pre-bleach images and an appropriate time-lapse and length for recovery (every 60 ms for 10 s for half and full FRAP of liquid droplets). Images were expanded with bilinear interpolation for optimal visualization.

For half FRAP analysis, each recovery curve was collected, then normalized to minimum and maximum. Then a group of at least 19 curves were averaged together and fit with a two component exponential recovery function. 5 replicates of this process yielded 5 sets of fit components that were then averaged together (n = 141 total) to yield the final results. For full FRAP, each curve was fit with a one component exponential recovery function. All fits were averaged together to yield the final results (n = 28). Monomer control FRAP was treated the same way as full FRAP (n = 70).

Sphericity analyses were limited to thresholded puncta with sizes > 350 nm. The aspect ratio is the ratio of the lengths of the long axis and short axis.

ThT localization assays were performed on an LSM 780 microscope in photon counting lambda mode with a 100x alpha Plan-Apochromat (NA = 1.46) objective. Excitation of ThT and mEOS was achieved at 405nm. Spectra for ThT and mEOS were obtained empirically from the data and used to linearly unmix the images. Unmixed images were then smoothed with a 1.5 pixel standard deviation Gaussian blur, cropped and the contrast was adjusted for optimal viewing of the signal in the punctate regions.

### Sup35 colony color assay

Photoconverted yeast cells were sorted 750 events each from high and low AmFRET populations by a BD Influx Sorter each into 200µl of SD CSM. Subsequently, the media containing sorted cells were spread with glass beads onto ¼ YPD plates; these plates enhance visualization of color effects subsequent to adenine deficiency for the Sup35-C red/white colony assay (Alberti et al., 2009). After growth at 30°C for 2–3 days, plates were moved to 4°C overnight to deepen the red coloration, and photographed the following day.

### FACS sorting of Sup35 PrD #24 expressing cells

Photoconverted yeast cells were sorted 500,000 events each from high and low AmFRET gates (using Sup35 PrD of [*PIN+*] cells and unfused mEos3.1 cells as positive and negative readouts of AmFRET, respectively) in a BD Influx Sorter, each into 1 ml of SGal-Ura. The cells were then concentrated by centrifugation and used for half-FRAP immediately.

### Semi-Denaturating Detergent Agarose Gel Electrophoresis (SDD-AGE)

The SDD-AGE procedure was adapted from (Halfmann and Lindquist, 2008) were lysed by bead-beating with a 2010 Geno/Grinder^®^, sarkosyl was used in the sample buffer, and the gel was directly imaged with GE Typhoon™ Imaging System. Images were then background subtracted using a 250 pixel rolling ball, cropped and contrast-adjusted.

---

**Table S1. Parameters of fits of DAmFRET datasets**

**Table S2. Plasmids used and polypeptide sequences characterized in this study**

**Table S3. Yeast strains used in this study**

**Supplementary Figure 1, related to Fig. 1.**

(A) Autofluorescence accurately distinguishes live from dead cells. Yeast cells expressing mEos3.1 (not photoconverted) were stained with Sytox Far Red. A histogram of Sytox intensity for single cells is shown on the left, and the Sytox-positive population (“dead”) is colored red in the dot plot of blue autofluorescence (405 nm excitation, 457/45 nm emission) vs. mEos3.1 (donor) intensity on the right. Based on this relationship, DAmFRET experiments excluded the population with blue autofluorescence and lacking detectable mEos3.1 fluorescence.

(B) The indicated proteins were expressed in isogenic [*psi-*] [*rnq-*] or [*PSI+*] [*RNQ+*] strains and tested for donor intensity prior to photoconversion and then for acceptor intensity following photoconversion. Samples containing amyloid are indicated in red; non-amyloid structured assemblies in blue; and unstructured or monomer in black. The linear regression (R^2^ = 0.9728) indicates that mEos3.1 photoconversion is independent of fusion, expression level, or structure of the mEos3.1-tagged query protein.

(C) Density plot showing tight correlation of donor and acceptor fluorescence of mEos3.1 following the photoconversion step.

(D) Left-right: A sample image of Spc42-mEos3.1 from the ImageStream^®x^ MkII, a 2D density histogram showing the relationship between Spc42 spot size and peak amplitude showing the selected population for analysis, and a plot of the peak amplitude of beads on the ImageStream as a function of objective focus Z Shift showing the shape of the focal volume (See Methods).

(E) Left: histograms of Spc42-mEos3.1 peak amplitude at different laser powers and Right: the trend of the centers of these histogram peaks (See Methods).

(F) Representative images of High-FRET cells from DAmFRET data of ASC. The panels are (left to right) donor, FRET, acceptor, and bright field merged with acceptor signal.

(G) DAmFRET plot of cells expressing mEos3.1 without a fusion partner.

**Supplementary Figure 2, related to Fig. 2.**

Corresponding DAmFRET plots of the curve fits shown in Fig. 2D.

**Supplementary Figure 3, related to Fig. 3.**

(A) DAmFRET plots of known yeast prion-forming sequences and of Ngr1 PrL and Sla1 PrL.

(B) Images of cells acquired by imaging flow cytometry. The cells were selected from high or low-AmFRET populations within the same concentration range, expressing the indicated protein from (A). Representative images for each population are shown. BF, Bright Field.

(C) Representative images showing ThT-stained cells expressing Sla1 PrL or Ngr1 PrL.

(D) DAmFRET of the PrD of a well-characterized fungal prion HET-s and its known hypomorphic mutant W287A. The inset shows **δ** ± error. *The **δ** value for W287A is likely underestimated due to the Weibull function necessitating that the high AmFRET fraction goes to 1.

**Supplementary Figure 4, related to Fig. 4.**

(A) DAmFRET plots and the indicated gates used for the quantification of mean AmFRET shown in Fig. 4A.

(B) Intensity (arbitrary units) of the two puncta indicated by arrows in Fig. 4E immediately before (red and blue bar) and after (purple bar) coalescing with each other.

(C) Bar graphs depicting the quantification of FCS, with the molecular brightness of GFP in cells expressing 1x GFP or endogenous, full-length Sup35 fused to GFP. See Methods for details.

**Supplementary Figure 5, related to Fig. 5.**

(A) DAmFRET plots of Sup35 PrD mutants and scrambled variants, as an extension of those shown in Fig. 5A.

(B) DAmFRET of Sup35 PrD^Q^ in cells with different endogenous amyloid templates ([*PIN*+], [*PSI+*^*weak*^], or [*PSI+*^*strong*^]) in comparison with that of Sup35 PrD WT. Mean AmFRET values are denoted as µ.

(C) Representative confocal images depicting ThT staining in cells expressing Sup35 PrD^Q^ or Sup35 PrD #24. (D) SDD-AGE of Sup35 PrD #24 and Sup35 PrD^Q^ expressed in [*PIN+*] or [*pin-*] cells, along with WT Sup35 PrD as a positive and negative control for amyloid.

